# Predicting leaf traits across functional groups using reflectance spectroscopy

**DOI:** 10.1101/2022.07.01.498461

**Authors:** Shan Kothari, Rosalie Beauchamp-Rioux, Florence Blanchard, Anna L. Crofts, Alizée Girard, Xavier Guilbeault-Mayers, Paul W. Hacker, Juliana Pardo, Anna K. Schweiger, Sabrina Demers-Thibeault, Anne Bruneau, Nicholas C. Coops, Margaret Kalacska, Mark Vellend, Etienne Laliberté

## Abstract

- Plant ecologists use functional traits to describe how plants respond to and influence their environment. Reflectance spectroscopy can provide rapid, non-destructive estimates of leaf traits, but it remains unclear whether general trait-spectra models can yield accurate estimates across functional groups and ecosystems.
- We measured leaf spectra and 22 structural and chemical traits for nearly 2000 samples from 104 species. These samples span a large share of known trait variation and represent several functional groups and ecosystems. We used partial least-squares regression (PLSR) to build empirical models for estimating traits from spectra.
- Within the dataset, our PLSR models predicted traits like leaf mass per area (LMA) and leaf dry matter content (LDMC) with high accuracy (*R^2^*>0.85; %RMSE<10). Models for most chemical traits, including pigments, carbon fractions, and major nutrients, showed intermediate accuracy (*R^2^*=0.55-0.85; %RMSE=12.7-19.1). Micronutrients such as Cu and Fe showed the poorest accuracy. In validation on external datasets, models for traits like LMA and LDMC performed relatively well, while carbon fractions showed steep declines in accuracy.
- We provide models that produce fast, reliable estimates of several widely used functional traits from leaf reflectance spectra. Our results reinforce the potential uses of spectroscopy in monitoring plant function around the world.

## Introduction

Plant functional ecology relies on the measurement of traits that influence how plants interact with their abiotic and biotic environment. Such traits can be used to interpret or predict outcomes at the community- or ecosystem-scale (Funk et al. 2017), ranging from species sorting across environmental gradients (Kunstler et al. 2016) to rates of carbon and nutrient cycling (Cornwell et al. 2008; Ollinger et al. 2008). During the last 20 years, ecologists have developed standardized trait protocols and databases to enable broad comparisons of plant functional strategies across species (Perez-Harguindeguy et al 2013; Kattge et al. 2020). Nevertheless, because trait measurement campaigns can be costly and challenging, trait databases remain sparse and much of the world’s plant diversity is poorly represented (Kattge et al. 2020). The resources needed to carry out intensive trait measurement campaigns using conventional methods has motivated the search for other approaches that could yield fast, reliable trait estimates.

One such approach is to predict plant traits from reflectance spectra, which report the fraction of light reflected from a surface across a range of wavelengths. For vegetation, the range most often studied is around 350-2500 nm, which includes >97% of solar radiation (American Society for Testing and Materials 2020). Because the structure and chemistry of plant tissues determine their optical properties, many traits can be estimated from reflectance spectra measured at leaf or canopy scales (Jacquemoud & Ustin 2019). At the leaf level, linking reflectance spectra and functional traits provides a fast way to estimate large numbers of leaf traits from individual plants. At the canopy level, it enables researchers to map traits using remote sensing, which may enable monitoring of plant community structure (Durán et al. 2019; Zheng et al. 2021) or biogeochemical cycles (Wessman et al. 1988; Chadwick & Asner 2018) over entire landscapes. Understanding leaf spectral variation is essential for scaling up to the canopy level (Asner et al. 2011), and leaf-level spectral estimates of traits are part of many workflows for canopy trait mapping (Wang et al. 2020). These uses of leaf spectra make it important to understand their functional drivers (Kothari & Schweiger 2022).

Methods to predict traits from leaf or canopy spectra can be summarized into three groups: (1) simple indices calculated from reflectance at a few wavelength bands (e.g. Sims & Gamon 2002); (2) physics-based radiative transfer models (Jacquemoud et al. 2009; Féret et al. 2017); and (3) multivariate empirical methods that use large portions of the spectrum to build trait prediction models. Among this last group of methods, partial least-squares regression (PLSR) is the most common, although others like stepwise regression and (more recently) deep learning approaches like convolutional neural networks are also used. These methods are less mechanistic but offer the flexibility often needed to predict traits associated with complex optical properties (Curran 1989). For example, predicting leaf nitrogen concentrations using radiative transfer models has been a persistent challenge because leaves contain nitrogen in many forms, including proteins (e.g. RuBisCO) and pigments (Wang et al. 2018; Féret et al. 2021). Moreover, the absorption features of nitrogen-containing bonds often overlap with others, especially after being broadened by multiple scattering (Curran 1989). As a result, leaf nitrogen is not identifiable with unique, distinct absorption features, and is often best predicted using empirical approaches like PLSR (Asner et al. 2011; Yan et al. 2021). Researchers have likewise built PLSR models to estimate a wide variety of foliar traits from fresh-leaf spectra, including pigments (Yang et al. 2016), non-structural carbohydrates (Ely et al. 2019), and condensed tannins (Couture et al. 2016).

Estimating traits from leaf spectra has several advantages that may compensate for the sparsity of trait datasets. Compared to measuring a large number of traits, measuring spectra is fast, has low marginal cost, can be non-destructive, and requires relatively little training. But since PLSR is purely empirical rather than mechanistic, a model calibrated using one dataset may yield inaccurate or biased predictions on another dataset if the association between traits and optical properties varies among contexts (Yang et al. 2016; Wang et al. 2020). The problem may be most severe when the model relies on absorption features that are not causally associated with the target trait, but rather with other covarying traits (“constellation effects” *sensu* Chadwick & Asner 2016; Nunes et al. 2017). This situation may occur most often when the target trait lacks major, well-defined absorption features, like concentrations of many elements (Nunes et al. 2017). If trait covariance patterns vary among datasets, overfitting to the training data could cause such models to produce inaccurate trait estimates from new spectral data (Kothari & Schweiger 2022). Empirical models may be especially fallible beyond the range of data used to train them (Schweiger 2020; Burnett et al. 2021).

To enable the widespread use of spectral models for trait prediction, it is essential to build general models that encompass the broadest possible range of spectral and trait diversity (Serbin et al. 2019; Wang et al. 2020). Such models could allow researchers to estimate plant traits from spectra in their study systems without going through the laborious process of building their own models. Asner et al. (2011) invoked this goal in developing spectral models for many leaf traits in humid tropical forest canopies around the world. In contrast, Serbin et al. (2019) focused on a single trait (leaf mass per area; LMA) but included several functional groups and biomes from the tropics to the tundra. We sought to push this goal further by including both a large number of leaf traits and a broad range of ecosystems and functional groups, allowing us to compare which traits are the most practical targets for estimation by general spectral models.

We compiled foliar traits and spectra measured using standardized protocols on nearly 2000 samples that include trees, shrubs, graminoids, and other herbs from a range of ecosystems (Table 1). We included structural and biochemical traits that relate to several aspects of leaf function, including nutrient economics, water relations, photosynthesis and photoprotection, and structural defense. We focused on building models to predict these traits from reflectance spectra, but we also compared the performance of reflectance, transmittance, and absorptance spectra. Among these properties, reflectance is the simplest and most common to measure because it bears the most relevance for interpreting remotely sensed canopy reflectance and does not require an integrating sphere. But since the content of leaf chemical constituents most directly alters the amount of light absorbed (Jacquemoud & Ustin 2019), we conjectured that absorptance spectra might be better for estimating traits, and aimed to determine whether any potential benefit transmittance or absorptance spectra provide in trait estimation outweighs their greater costs in time and equipment.

**Table 1:** A summary of all projects with samples included in the study.

| Project | Description | Latitude range | Longitude range | Species | Samples per functional group |  |  |  |  |  |  | Total |
| --- | --- | --- | --- | --- | --- | --- | --- | --- | --- | --- | --- | --- |
|  |  |  |  |  | Broadleaf | Conifer | Graminoid | Fern | Forb | Shrub | Vine |  |
| Beauchamp-Rioux | Nine common broadleaf tree species measured from June to October 2018 at multiple forested sites in Québec (Beauchamp-Rioux 2022) | 45.4-46.0 N | 73.3-75.7 W | 9 | 436 |  |  |  |  |  |  | 436 |
| Blanchard | A sampling of nearly all tree species found in southern Québec, collected across multiple sites throughout summer 2019 | 45.1-46.0 N | 71.1-74.0 W | 43 | 270 | 68 |  |  |  | 3 |  | 341 |
| Boucherville 2018 | A variety of plants sampled in August 2018 from the Parc national des Îles-de-Boucherville in southern Québec | 45.6-45.7 N | 73.5-73.6 W | 12 |  |  | 14 |  | 30 | 22 | 6 | 72 |
| Boucherville 2019 | A variety of plants sampled in July 2019 from the Parc national des Îles-de-Boucherville in southern Québec | 45.6 N | 73.5 W | 19 | 40 |  | 38 |  | 59 | 26 | 10 | 173 |
| CABO General | A small number of samples from southern Québec not collected as part of a specific project | 45.5-46.0 N | 71.9-74.1 W | 13 | 33 |  | 6 |  | 3 | 39 |  | 81 |
| Crofts | Trees sampled in July 2019 across an altitudinal gradient in southeastern Québec, along the transition from temperate deciduous to boreal forest | 45.4-45.5 N | 71.1-71.2 W | 25 | 139 | 60 |  |  |  | 5 |  | 204 |
| Girard | Four common bog plants sampled in summer 2018 in Ontario and southern Québec across a gradient of nitrogen deposition (Girard et al. 2020) | 45.4-46.8 N | 71.1-75.5 W | 4 |  |  | 23 |  |  | 72 |  | 95 |
| Hacker | Plants sampled in May 2019 to evaluate functional trait variation throughout an endangered oak savannah in British Columbia (Hacker et al. 2022) | 48.8-49.3 N | 123.1-123.7 W | 23 | 10 |  | 38 | 7 | 90 | 56 |  | 201 |
| *Phragmites*<br>Temporal | Six co-occurring wetland plant species sampled from June to September 2019 in Boucherville, Québec (Pardo 2021) | 45.6 N | 73.4-73.5 W | 6 |  |  | 240 |  | 60 |  |  | 300 |
| Warren | *Agonis flexuosa* sampled in November 2018 along a chronosequence of increasing soil age in southwestern Australia (Turner et al. 2018) | 34.6 S | 115.9 E | 1 | 68 |  |  |  |  |  |  | 68 |
| Total |  |  |  | 104 | 996 | 128 | 359 | 7 | 242 | 223 | 16 | 1971 |

The performance of statistical models is often evaluated by splitting a dataset at random into calibration and validation subsets. This procedure represents a best-case scenario, since the data are all collected with the same protocols and, based on the mathematics of random sampling, the two subsets must have similar distributions of predictor and response variables. Therefore, we validated our models on both a random subset of our dataset (internal validation) and on independent datasets (external validation)—the latter perhaps representing a more realistic test of how they would perform in practical applications.

Concerns about the generality of spectral trait models also raise the question: What factors determine whether a trait model yields accurate predictions on an external dataset? Many existing models to predict leaf traits were built using samples belonging to a single functional group or ecosystem, leaving it an open question how they would transfer to new settings. One might expect that models would show poorer performance as the calibration samples (used to build the model) and the external dataset’s samples become less functionally similar (Yang et al. 2016). But if the models show consistently strong performance, it would suggest that associations between traits and optical properties are robust, not contingent on the particular samples included. To examine model transferability, we divided our dataset by functional group and examined model accuracy using each functional group in turn as the validation dataset for models calibrated using the remaining functional groups. We posed two hypotheses about these analyses: (1) For a given trait, predictions are most accurate when the distributions for that trait are most similar between the calibration and validation datasets; and (2) traits whose predictions are most accurate across the full dataset will also yield the most accurate predictions (on average) when transferring models onto left-out functional groups. We aimed to shed light on the factors that influence a model’s accuracy across a range of settings.

## Methods

### Sampling

We assembled a database (*n* = 1971; species = 104) of leaf spectra and functional traits measured as part of the Canadian Airborne Biodiversity Observatory (CABO; www.caboscience.org). We aggregated data from ten individual projects carried out during 2018 and 2019 at sites across temperate Canada and one site in southwestern Australia (Table 1). The sites span a wide range of edaphic properties, from nutrient-poor bogs and phosphorus-limited woodlands to highly fertile post-agricultural and wetland sites. Each sample comprised a large, homogeneous group of sunlit leaves, which we divided up for spectral and trait measurements. We classified samples into seven functional groups: broadleaf trees, coniferous trees, ferns, forbs, graminoids, shrubs, and vines (Table 1). Further details on sampling protocols and functional groups are found in the Supplementary Materials.

### Leaf spectral and trait measurements

We measured directional-hemispherical reflectance and transmittance spectra (350-2500 nm) using an HR-1024i spectroradiometer equipped with a DC-R/T integrating sphere from Spectra Vista Corporation (Poughkeepsie, NY, USA). For each sample, we measured spectra from the adaxial surface of six leaves or (for small or narrow leaves) leaf arrays. We resampled spectra to 1 nm resolution, averaged spectra from the same sample, and trimmed them to 400-2400 nm by removing the ends, which are often noisy. Finally, we calculated absorptance at each wavelength by subtracting the sum of reflectance and transmittance from 1. Further details on spectral measurement and processing can be found in the Supplementary Materials.

We measured the following traits on each leaf sample: Leaf mass per area (LMA), leaf dry matter content (LDMC), equivalent water thickness (EWT), and concentrations of soluble cell components, hemicellulose, cellulose, lignin, chlorophyll *a*, chlorophyll *b*, total carotenoids, and a number of elements (Al, C, Ca, Cu, Fe, K, Mg, Mn, N, Na, P, and Zn). Elements other than C and N were only available for a few projects that account for about one-third (*n* = 678) of the total samples. Further details on trait measurement protocols can be found in the Supplementary Materials. We compared our trait distributions to the TRY trait database to evaluate how well they span the global range of trait variation. Although it has known geographic and taxonomic biases, TRY is the most comprehensive database of its kind (Kattge et al. 2020). We also performed a principal components analysis (PCA) on the scaled and centered trait data to visualize patterns of trait variation among our samples.

Within a dataset, chemical traits may vary more in proportion to the mass or the area of the leaf (Osnas et al. 2013). When a trait is distributed mainly proportional to leaf area, mass-normalization induces a negative correlation with LMA. Since interspecific variation in pigments tends to be area-proportional, Kattenborn et al. (2019) used this principle, among others, to argue that pigment content (per unit area) is a better target than concentration (per unit mass) for remote sensing-based estimation. We applied our PLSR modeling approach to both mass- and area-based traits, but we focus on mass-based models since conventional protocols measure most chemical traits on a mass basis. Chemical traits should be assumed here to be mass-based when left unspecified. We also calculated the mass-proportionality of each chemical trait in our dataset as the ordinary least squares slope between the log_10_-transformed area-normalized trait and log_10_-transformed LMA (Osnas et al. 2013).

### PLSR modeling

#### Trait estimation

Due to the high dimensionality and multicollinearity of spectral data, we used a PLSR modeling approach to build trait estimation models. PLSR projects the high-dimensional input dataset (here, spectra) onto a smaller number of latent components constructed to maximize their covariance with the variable(s) to be predicted (traits). We built separate models using reflectance, transmittance, and absorptance as predictors, in each case using the whole trimmed spectrum (400-2400 nm). We did not transform the distributions of any trait to reduce skewness, as is sometimes recommended (Burnett et al. 2021); in preliminary tests, such transformations did not yield any consistent improvement in model performance. We performed all analyses in *R* version 3.6.3 (R Core Team 2020).

We calibrated and validated our main set of models (‘primary models’) largely following the example of Serbin et al. (2014) and Burnett et al. (2021) using *R* package *pls* v. 2.7.1 (Mevik et al. 2019). First, we divided the full dataset into 75% calibration and 25% validation sets at random, stratified by functional group (see Table 1). We fit a preliminary PLSR model for each trait on the calibration subset, using 10-fold cross-validation to select the number of components to use in our final models. We chose the smallest number of components that brought the cross-validation root mean squared error of prediction (RMSEP) within one standard deviation of the global minimum—a distinct number (4-27) for each trait and type of spectrum. We calculated the variable influence on projection (VIP) metric (Wold et al. 2001) to determine which spectral regions were most important for predicting each trait.

Next, we fit our final models through a jackknife resampling procedure in which we further divided the 75% calibration dataset 100 times into random 70% testing and 30% training subsets. For each iteration, we built a model from the 70% testing dataset using the previously chosen number of components and applied it to the 30% training dataset. This procedure allowed us to get a distribution of summary statistics that shows how model performance varies based on random changes in the training and testing datasets. The primary model comprises the 100 sets of coefficients for each trait—one from each resampling iteration—which we applied to the 25% validation dataset to get a distribution of estimates for each sample. We compared the measured values to the mean estimates for each trait to produce summary statistics (*R^2^*, RMSE, %RMSE) for internal validations. In all cases, we report RMSE relative to the 1:1 line. For robustness to outliers, we defined %RMSE as the RMSE divided by the range of the inner 95% of trait values, in contrast to a more common definition (Burnett et al. 2021) that uses the full range.

#### External validation

We compiled data from three sources as external validation datasets for our primary models. The first two are LOPEX (Hosgood et al. 1994) and ANGERS (Féret et al. 2008), which have been widely used to validate methods for plant trait estimation from reflectance spectra. Between them, these datasets include a very wide variety of temperate plants, many cultivated. The third source is an expanded version of the Dessain dataset previously described in Kothari et al. (2022), which includes herbs and trees from southern Québec. Unlike LOPEX and ANGERS, most of the Dessain dataset’s trait data were measured using the same protocols as the core CABO datasets. All three datasets were measured using integrating spheres and include similar spectral ranges, but the instruments and their specifications differ from each other and the core datasets we used to train the models, which can cause slight but important differences in measured reflectance (Lukeš et al. 2017; Hovi et al. 2018).

Since we did not measure transmittance spectra for the Dessain project, we focused on reflectance spectra from the validation datasets. We applied our sets of reflectance-based coefficients from resampling analyses (100× for each trait) to the spectra in each dataset and took the mean of the resulting 100 estimates for each sample for comparison to the measured traits. Not all traits were available for each dataset. Further details on our handling of external validation data are in the Supplementary Materials.

#### Model transferability among functional groups

Besides assessing the viability of our primary models, we also aimed to test whether and when models could estimate traits accurately on a group of species functionally distinct from those used in calibration (Hypotheses 1 and 2). To this end, we built another set of models: we selected a single functional group at a time as a validation dataset and used the reflectance spectra of the remaining functional groups to calibrate a PLSR model for each trait—using 10-fold cross-validation and component selection as above, but omitting the jackknife procedure. Finally, we applied the model to the validation functional group. We repeated this procedure using each functional group in turn as the validation dataset, besides ferns and vines because of their low numbers of samples. Among elements, we only considered C and N because other elements were only available for certain projects and we could not guarantee adequate coverage of all functional groups. We quantified performance using *R^2^*, RMSE, and %RMSE—calculating %RMSE using the inner 95% trait range of the full dataset in the denominator rather than the inner 95% trait range of only the validation functional group. This metric, which we call %RMSE_full_ for clarity, avoids apparent reductions in model performance due to a narrow trait range in the validation functional group.

We conjectured that for a given trait, model performance would decline as the calibration and validation datasets became more dissimilar in their trait distributions (Hypothesis 1). We quantified dissimilarity using the Hellinger distance between the calibration and validation datasets’ distributions for each trait, as estimated in *R* package *statip v. 0.2.3* (Poncet et al. 2019). Here we focused on %RMSE_full_ because it is normalized to the scale of the data (unlike RMSE) and incorporates bias (unlike *R^2^*). We tested the influence of trait dissimilarity on %RMSE_full_ with analysis of covariance (ANCOVA) using Hellinger distance (continuous), the trait identity (categorical), and their interaction as predictors.

## Results

### Trait and spectral variability

The ranges for most traits in our dataset spanned most of the trait ranges in TRY. Many traits, including LMA, EWT, and all elements other than C and N, had distributions with strong positive skew in both TRY and CABO data, resulting in a long upper tail of high values (Fig. 1). For certain elements (Al, Cu, Ca, Fe, K, Mg), the CABO dataset had less representation of values along the upper tail than TRY. For other elements (Na, P, Zn), as well as LDMC, N, and pigments, the CABO data had greater representation of high values. For yet other traits (LMA and C), the CABO data had a somewhat narrower range towards either extreme.

**Fig. 1:**
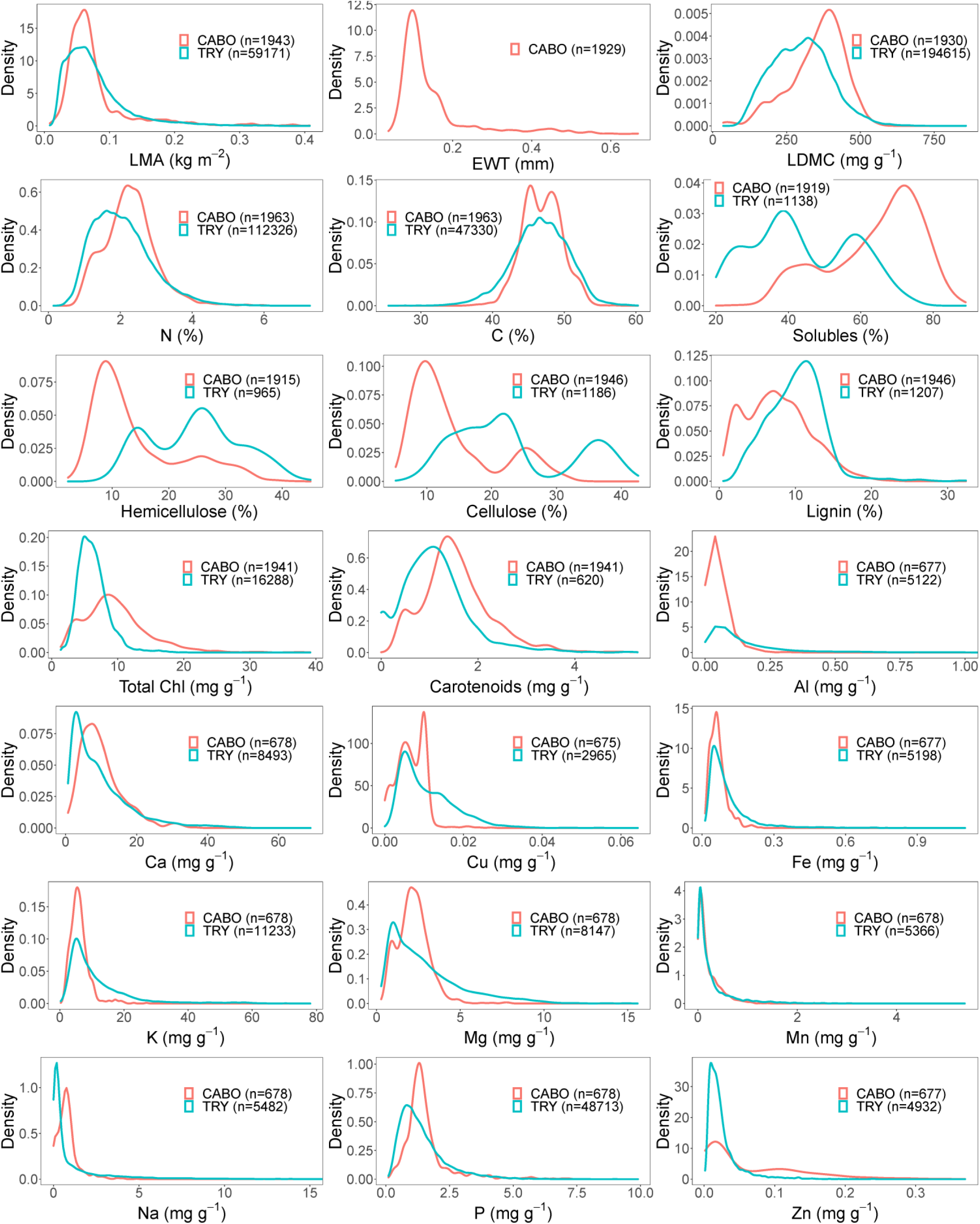
Density plots comparing the distributions of various leaf structural and chemical traits in the CABO dataset to the TRY database (Kattge et al. 2020). We filtered TRY data to include only values with an error risk below 3. EWT is only shown for CABO because it is not a trait in TRY. Elements besides C and N are only available for the Beauchamp-Rioux, Boucherville 2018, Girard, and Warren projects.

Within the dataset, functional groups often differed in their trait values, as shown by a principal components analysis (PCA) of the centered and scaled trait data (excluding elements other than C and N; Fig. S1). The first two PCs of the trait data explained 65.7% of total variance. Graminoids were distinguished by high concentrations of hemicellulose and cellulose and low concentrations of soluble carbon, lignin, and total C. Leaf mass per area and LDMC were negatively correlated with N and pigment concentrations; conifers (besides deciduous *Larix laricina* (Du Roi) K. Koch) and the evergreen broadleaf tree *Agonis flexuosa* (Willd.) Sweet had particularly high LMA and LDMC, while other functional groups each contained significant variation along this axis. High values of EWT (>0.3 mm) were dominated by evergreen conifers and graminoids adapted to wetlands, such as *Typha angustifolia* L. and *Phragmites australis* (Cav.) Trin. ex Steud. *A. flexuosa* represented most of the samples with the lowest concentrations of P, K, and Fe and the highest concentrations of Na, while bog shrubs and sedges had particularly low Ca concentrations. The large fraction of variance explained by major trait axes indicates that certain clusters of traits covary strongly, often in ways that correspond to functional groups.

Our samples showed the typical shape of leaf spectra, including a sharp increase in reflectance and transmittance (‘red edge’) between the visible and near-infrared (NIR; 700-1000 nm) ranges and prominent water absorption bands in the short-wave infrared (SWIR; >1000 nm) range (Fig. 2; Fig. S2). Compared to other functional groups, needleleaf conifers had low reflectance and transmittance and high absorptance across the spectrum. Forbs also tended to have lower transmittance and higher absorptance in the SWIR. In absolute terms, absorptance and transmittance spectra had greater variation than reflectance spectra, especially in the SWIR.

**Fig. 2:**
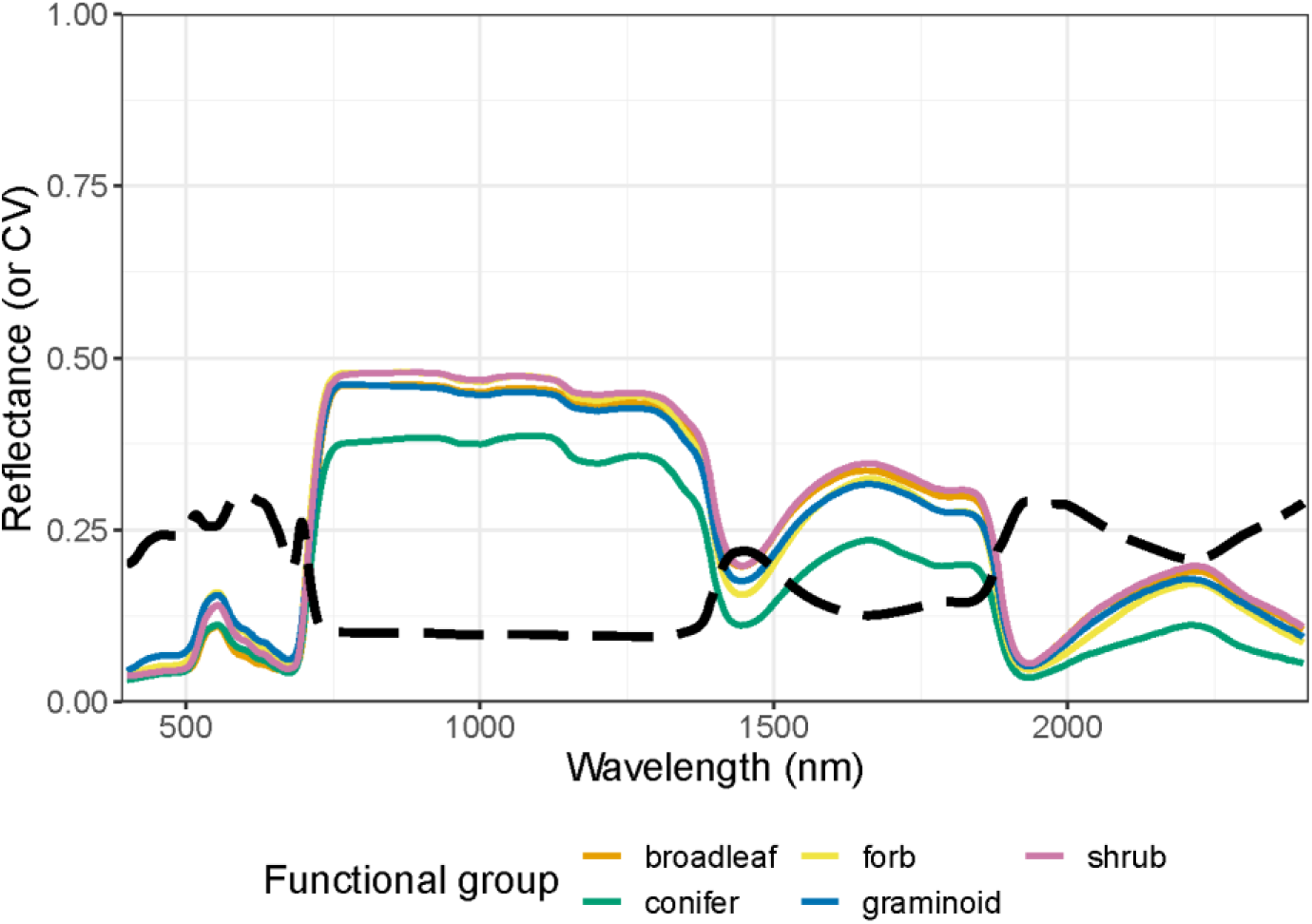
The median leaf-level reflectance at each wavelength across the spectrum, separated by functional group. We omit ferns and vines, which are only represented by one species each. The dashed line shows the coefficient of variation across all samples.

### PLSR modeling

#### Internal calibration and validation

The accuracy of trait estimates from reflectance spectra varied widely by trait (Table 2; Fig. 3-4; Fig. S3-9). Our primary models estimated structural and water-related traits—LMA (*R^2^*= 0.892; %RMSE = 8.04), EWT (*R^2^* = 0.885; %RMSE = 8.46), and LDMC (*R^2^* = 0.867; %RMSE = 9.78)—with high accuracy. Models for certain chemical traits such as carbon fractions (*R^2^*= 0.558-0.826; %RMSE = 12.7-19.1), major elements like C, N, K, and Ca (*R^2^* = 0.541-0.747; %RMSE = 13.2-20.4), and pigments (*R^2^* = 0.644-0.689; %RMSE = 13.8-15.0) also showed intermediate-to-high accuracy. Finally, models for some micronutrients like Cu (*R^2^* = 0.295; %RMSE = 25.9) or Mn (*R^2^* = 0.254; %RMSE = 22.8) performed very poorly. For pigments, N, and some other elements, models underestimated trait values at the sparsely represented upper tail of the measured range, while for C the models overestimated values at the sparse lower tail (Fig. 3-4). Trait models with lower accuracy on the internal validation data also had greater variability in accuracy across iterations in the resampling analysis—even when leaving out elements other than C and N, which are only available for a few projects (Fig. S10-12).

**Fig. 3:**
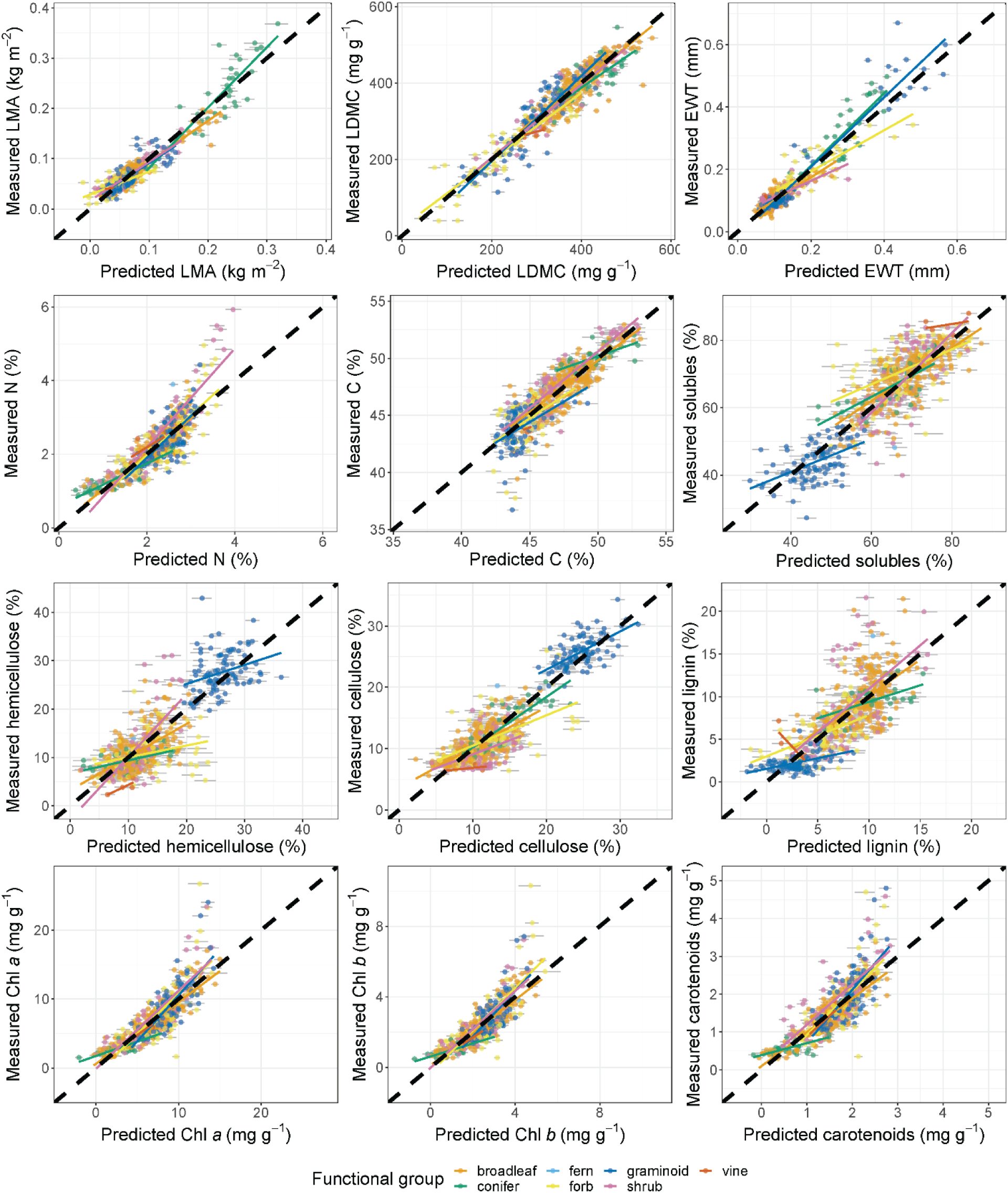
Plots of observations against reflectance-based PLSR predictions among internal validation data for various leaf structural and chemical traits. The black dashed line in each panel is the 1:1 line. Colored lines represent best-fit lines from OLS regression for each functional group. Error bars around each point represent 95% confidence intervals based on the ensemble of models produced in the 100× jackknife analysis.

**Fig. 4:**
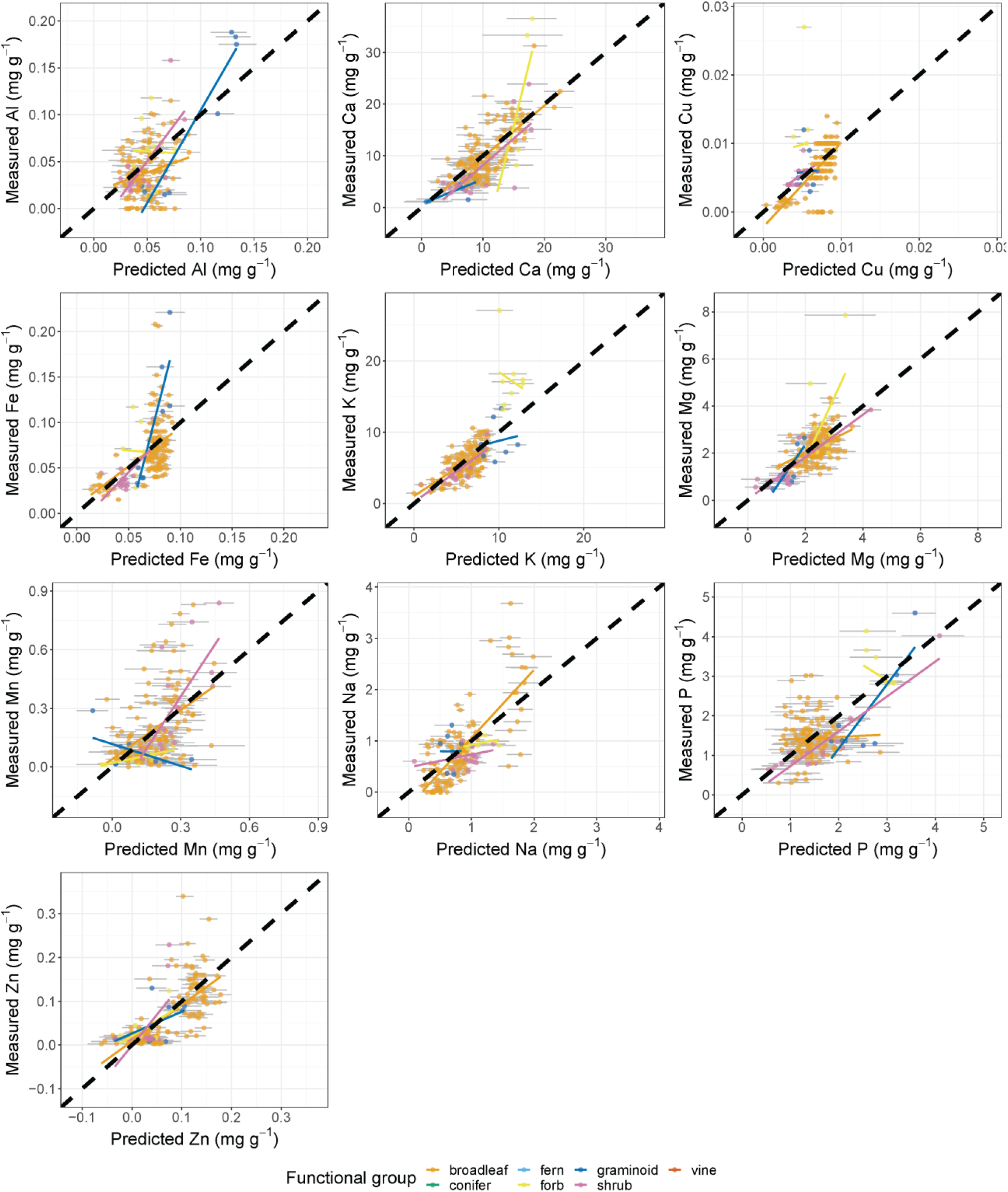
Plots of observations against reflectance-based PLSR predictions among internal validation data for elements other than C and N. These data are only available for the Beauchamp-Rioux, Boucherville 2018, Girard, and Warren projects. The black dashed line in each panel is the 1:1 line. Colored lines represent best-fit lines from OLS regression for each functional group. Error bars around each point represent 95% confidence intervals based on the ensemble of models produced in the 100× jackknife analysis.

**Table 2:** Summary statistics for the performance of reflectance-based models calibrated on CABO data and applied to internal and external validation datasets. %RMSE is calculated as RMSE divided by the inner 95% trait range. The column ‘# obs’ refers to the number of observations for the trait in the full CABO dataset.

|  | Internal |  |  |  |  | Dessain |  |  | LOPEX |  |  | ANGERS |  |  |
| --- | --- | --- | --- | --- | --- | --- | --- | --- | --- | --- | --- | --- | --- | --- |
| | # obs | # comps | $R^2$ | RMSE | %RMSE | $R^2$ | RMSE | %RMSE | $R^2$ | RMSE | %RMSE | $R^2$ | RMSE | %RMSE |
| LMA (kg m<sup>-2</sup>) | 1943 | 23 | 0.892 | 0.0175 | 8.04 | 0.593 | 0.0855 | 123 | 0.656 | 0.111 | 112 | 0.517 | 0.137 | 117 |
| LDMC (mg g<sup>-1</sup>) | 1930 | 20 | 0.867 | 32.9 | 9.78 | 0.824 | 40.6 | 13.6 | 0.860 | 46.4 | 12.0 |  |  |  |
| EWT (mm) | 1929 | 13 | 0.885 | 0.0334 | 8.46 | 0.629 | 0.0395 | 24.2 | 0.888 | 0.0705 | 27.5 | 0.825 | 0.0665 | 34.8 |
| N (%) | 1963 | 19 | 0.709 | 0.403 | 13.7 | 0.417 | 0.664 | 21.6 | 0.535 | 0.854 | 20.2 |  |  |  |
| C (%) | 1963 | 20 | 0.747 | 1.38 | 13.2 | 0.41 | 2.23 | 31.6 | 0.145 | 2.81 | 25.5 |  |  |  |
| Solubles (%) | 1919 | 23 | 0.733 | 6.59 | 15.1 | 0.284 | 10.9 | 26.7 |  |  |  |  |  |  |
| Hemicellulose (%) | 1915 | 20 | 0.695 | 4.35 | 16.1 | 0.324 | 5.28 | 19.7 |  |  |  |  |  |  |
| Cellulose (%) | 1946 | 25 | 0.826 | 2.74 | 12.7 | 0.327 | 6.68 | 43.5 | 0.353 | 5.47 | 23.1 |  |  |  |
| Lignin (%) | 1946 | 20 | 0.558 | 2.82 | 19.1 | 0.152 | 4.68 | 29.9 | 0.164 | 4.79 | 26.8 |  |  |  |
| Chl *a* (mg g<sup>-1</sup>) | 1941 | 13 | 0.689 | 2.07 | 13.8 | 0.312 | 2.88 | 26.7 | 0.409 | 2.72 | 28.1 | 0.503 | 2.80 | 21.9 |
| Chl *b* (mg g<sup>-1</sup>) | 1941 | 14 | 0.679 | 0.716 | 15.0 | 0.318 | 1.05 | 31.6 | 0.441 | 0.927 | 23.4 | 0.409 | 1.32 | 28.7 |
| Carotenoids (mg g<sup>-1</sup>) | 1941 | 12 | 0.644 | 0.434 | 14.7 | 0.195 | 0.585 | 27.0 | 0.288 | 0.694 | 24.8 | 0.419 | 1.09 | 29.9 |
| Al (mg g<sup>-1</sup>) | 677 | 6 | 0.304 | 0.0293 | 24.6 | 0.006 | 0.0813 | 33.5 |  |  |  |  |  |  |
| Ca (mg g<sup>-1</sup>) | 678 | 17 | 0.541 | 4.03 | 20.4 | 0.009 | 10.4 | 32.8 |  |  |  |  |  |  |
| Cu (mg g<sup>-1</sup>) | 675 | 5 | 0.295 | 0.00311 | 25.9 | 0.004 | 0.0147 | 34.0 |  |  |  |  |  |  |
| Fe (mg g<sup>-1</sup>) | 677 | 4 | 0.286 | 0.0295 | 23.0 | 0.045 | 0.053 | 26.6 |  |  |  |  |  |  |
| K (mg g<sup>-1</sup>) | 678 | 10 | 0.55 | 2.32 | 15.7 | 0.443 | 9.13 | 28.9 |  |  |  |  |  |  |
| Mg (mg g<sup>-1</sup>) | 678 | 18 | 0.394 | 0.714 | 22.8 | 0.149 | 2.3 | 53.6 |  |  |  |  |  |  |
| Mn (mg g<sup>-1</sup>) | 678 | 10 | 0.254 | 0.164 | 22.8 | 0.005 | 0.504 | 54.6 |  |  |  |  |  |  |
| Na (mg g<sup>-1</sup>) | 678 | 6 | 0.519 | 0.451 | 16.7 | 0.085 | 0.881 | 22.6 |  |  |  |  |  |  |
| P (mg g<sup>-1</sup>) | 678 | 15 | 0.313 | 0.607 | 20.5 | 0.051 | 2.06 | 51.2 |  |  |  |  |  |  |
| Zn (mg g<sup>-1</sup>) | 677 | 17 | 0.487 | 0.0486 | 24.2 | 0.045 | 0.118 | 34.9 |  |  |  |  |  |  |

The VIP metric revealed broad similarities in the regions of the spectrum that were important for predicting different traits from reflectance spectra (Fig. S13-15). In particular, major features of the visible region, including the green hump (centered at 550-560 nm) and especially the inflection point of the red edge (710-720 nm) were important for predicting most traits besides EWT. These visible-range importance peaks (especially at the red edge) were often higher for traits like Al, Zn, Mn, and hemicellulose than for pigments, which are the dominant drivers of visible reflectance (Jacquemoud & Ustin 2019). In general, VIP declined into the NIR but had several peaks in the SWIR, including relatively distinct peaks for multiple traits at 1390 nm and 1880 nm, and weaker ones at 1720 nm, 2020 nm, 2150 nm, and 2290 nm. For EWT, LMA, Cu, Fe, and Na, VIP remained high throughout a larger portion of the SWIR; uniquely, EWT had its global maximum VIP at 1390 nm.

In the dataset, most chemical traits were distributed mainly proportional to mass (Table S1). Exceptions included all pigments and several macro- and micronutrients, which showed mass-proportionality less than 0.5. We produced estimates of area-based chemical traits both by building models to estimate them directly and by multiplying estimated mass-based traits by estimated LMA (Table S1; Fig. S16-19). For most traits, estimating area-based traits directly yielded more accurate estimates; the difference was greatest for pigments, Ca, Cu, N, P, which all have area-proportionality less than 0.6. However, multiplying mass-based estimates by LMA estimates was slightly more accurate for many highly mass-proportional traits, including Al, C, Na, and cellulose.

For most traits, transmittance and absorptance spectra yielded slight improvements in performance relative to reflectance (Table S2; Fig. S3-12). Excluding traits that showed very poor performance in general (average *R^2^* < 0.35), the greatest improvements between reflectance and either transmittance or absorptance spectra (average Δ%RMSE > 1) occurred for LMA, LDMC, EWT, pigments, Ca, K, N, and P. Using transmittance or absorptance spectra never worsened performance considerably (average Δ%RMSE < -1).

#### External validation

We applied our reflectance-based primary models for each trait to three external validation datasets whose spectral and trait measurement protocols differed from the CABO dataset. In general, models performed less well on these independent datasets than the internal validation (Table 2; Fig. 5-6; Tables S3-4; Fig. S20). The traits that had the highest *R^2^* between observed and predicted values in the internal validation also had the highest *R^2^* in the external validation. As a result, structural and water-related traits (LMA, LDMC, and EWT) had moderate-to-high *R^2^* in the external validation (0.589-0.842 averaged across datasets). The RMSE for LDMC predictions was only slightly higher than in the internal validation (40.6-46.4 vs. 32.9 mg g^-1^). In contrast, both LMA and EWT were greatly overestimated for most samples, so even though the *R^2^* was moderate-to-high, RMSE was also high (LMA: 0.0855-0.137 vs 0.0175 kg m^-2^; EWT: 0.0395-0.0705 vs 0.0334 mm). Pigments, N, and particularly C and carbon fractions had correspondingly lower prediction accuracy and varying amounts of bias. Many other macro- and micronutrients (besides K) had very low prediction accuracy (*R^2^* < 0.15).

**Fig. 5:**
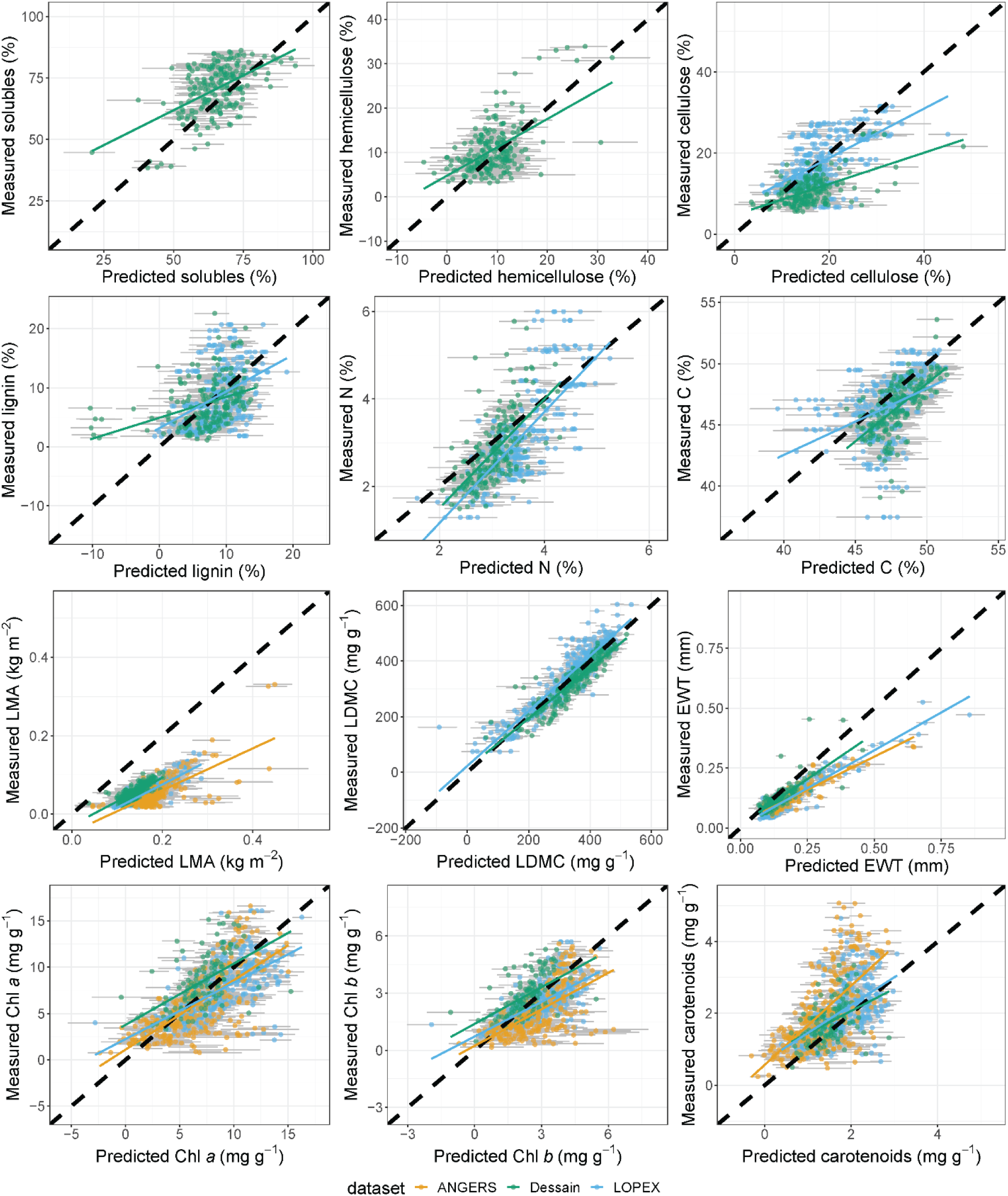
Plots of observations against reflectance-based PLSR predictions among external validation data for various leaf structural and chemical traits. The black dashed line is the 1:1 line. Colored lines represent best-fit lines from OLS regression for each dataset. Error bars around each point represent 95% confidence intervals based on the ensemble of models produced in the 100× jackknife analysis.

**Fig. 6:**
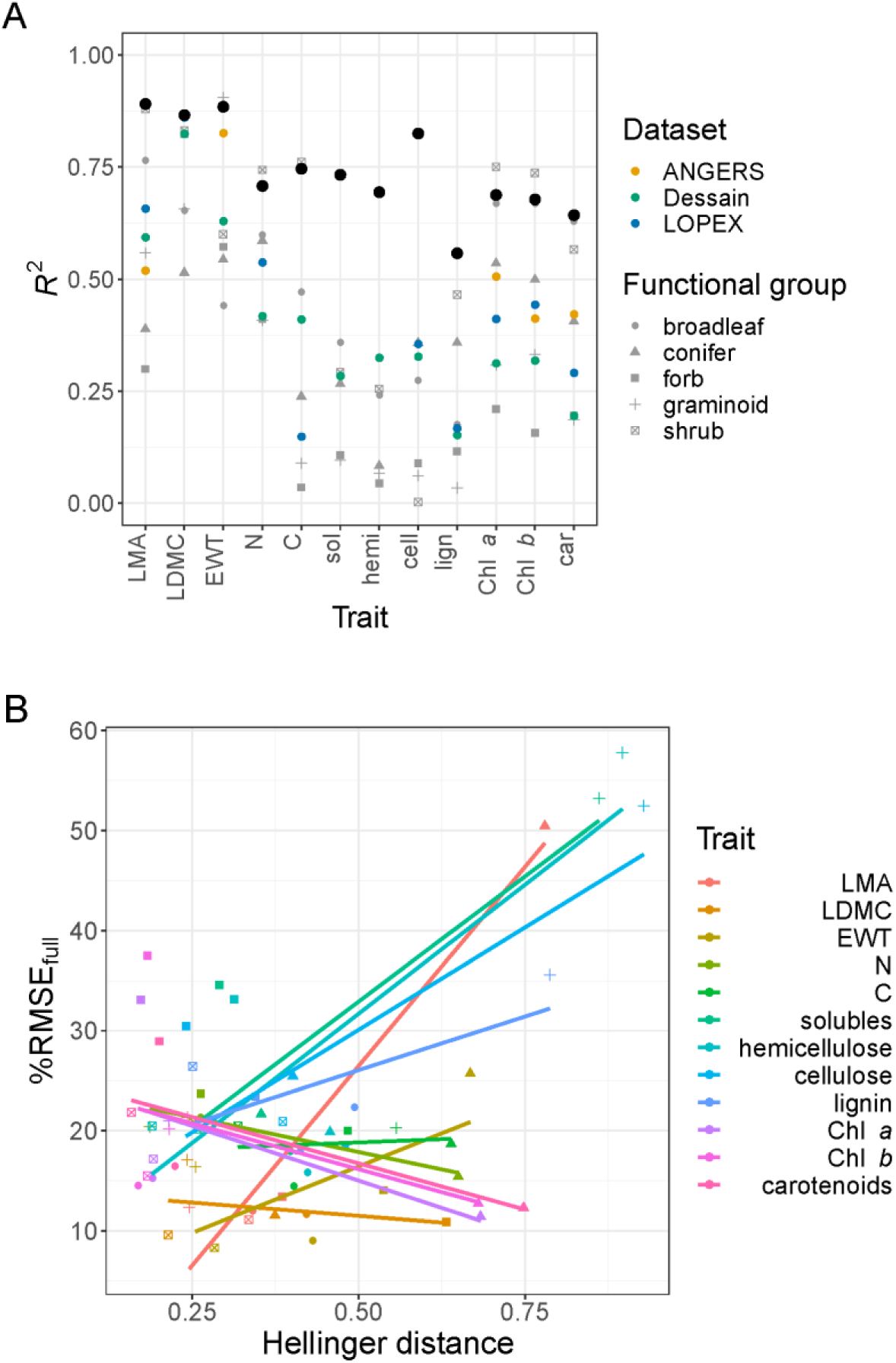
(A) A summary of model *R^2^* for each trait. Colors indicate whether the *R^2^* describes performance of models on: the random 25% internal validation dataset (black dots); specific functional groups used as validation data in model transferability analyses; or external validation datasets. (B) The relationship between %RMSE and Hellinger distance between calibration and validation trait distributions in model transferability analyses. Functional group labels are shared between panels and refer to the validation functional group.

#### Model transferability among functional groups

For each trait, models generally performed worse when applied to a functional group left out from the calibration dataset than when (as in our primary models) calibrated and applied to random selections of samples (Fig. 6-7; Table S5). In particular, models calibrated using the other functional groups tended to show poor performance when applied to graminoids or forbs. Carbon fractions tended to be estimated with relatively low accuracy (high RMSE, low *R^2^*); because these carbon fractions were predicted well within the internal validation, this result weakened the relationship posited in Hypothesis 2 between average model performance in internal validation and in functional group transferability analyses. Models for carbon fractions applied to graminoids yielded particularly biased predictions, with %RMSE_full_ > 50 for all but lignin. While graminoids were distinctively high in hemicellulose and cellulose and low in solubles and lignin, the model predicted them to be much more similar to the other functional groups (Fig. 7). Models for LMA applied to conifers likewise showed %RMSE_full_ > 50 because the distinctively high values of evergreen (non-*Larix*) conifers were underestimated.

**Fig. 7:**
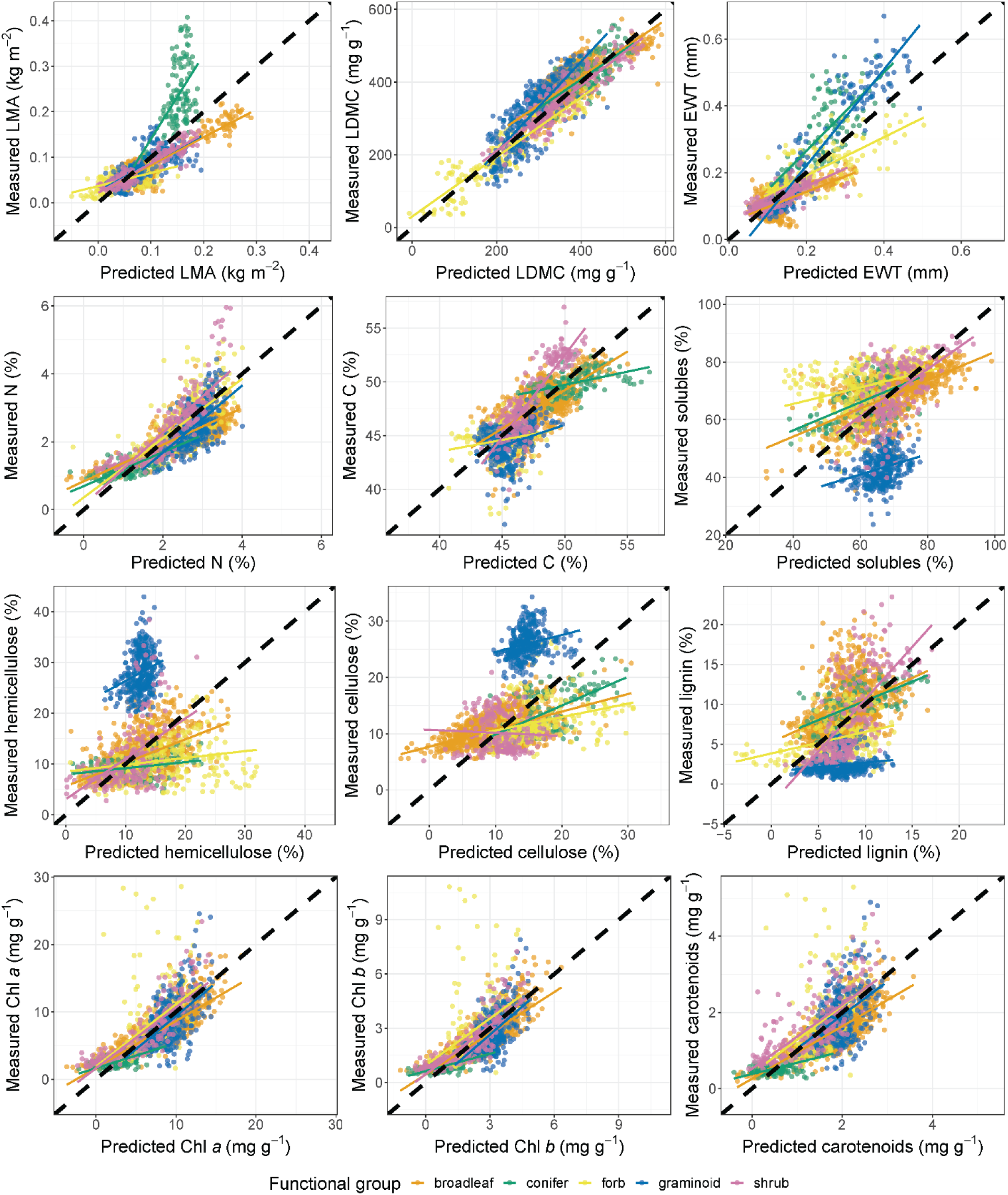
Plots of observations against reflectance-based PLSR predictions from model transferability analyses for various leaf chemical and structural traits. Here, we estimated traits for samples of each functional group from models calibrated on all the remaining functional groups—within panels, predictions for each functional group come from distinct models. The black dashed line in each panel is the 1:1 line. Colored lines represent best-fit lines from OLS regression for each functional group.

In our ANCOVA model, there were significant effects of trait (*F*_11,36_ = 2.626; *p* = 0.014), Hellinger distance (*F*_1,36_ = 21.066; *p* < 10^-4^), and their interaction (*F*_11,36_ = 4.139; *p* < 0.001) on %RMSE_full_. This result led us to perform separate linear regressions between %RMSE_full_ and Hellinger distance for each trait (Fig. 6B). The relationship was significant and positive for LMA (*t*(3) = 7.141; *p* = 0.006), but not significant and highly variable in strength and direction among the other traits. Support for Hypothesis 1 was thus also weak and equivocal.

## Discussion

We built PLSR models to predict widely used foliar traits from reflectance spectra across nearly 2000 samples of 104 species, including several functional groups and ecosystems. Our findings underscore that leaf spectra integrate many aspects of leaf phenotypes in a single measurement (Cavender-Bares et al. 2017; Kothari & Schweiger 2022) and help explain why spectral variation can serve as a surrogate for phenotypic variation (Schweiger et al. 2018; Frye et al. 2021).

### Comparing PLSR model performance among traits

The performance of our primary models was highest for structural and water-related traits (LMA, LDMC, and EWT), corroborating studies that have reported high accuracy using diverse modeling approaches (Serbin et al. 2019; Guzmán Q. et al. 2021). Indeed, the contents of dry matter and water are strong drivers of optical properties across much of the NIR and SWIR (Féret et al. 2008). Our models predicted N, C, individual carbon fractions, and pigments with intermediate accuracy. These traits also have known influences on leaf optical properties. Pigments control optical properties in the visible range (Jacquemoud & Ustin 2019) and correlate with N. Cellulose, lignin, and N-containing compounds like proteins also have bonds with known (if weak) absorption features, primarily in the NIR and SWIR (Curran 1989).

Models for elements at low concentrations often showed poor performance. Most of these elements have no measurable absorption features in this range, so their concentrations can only be estimated indirectly, often through symptoms of nutrient deficiency (Jacquemoud & Ustin 2019). In some cases, our models could predict variation across, but not within, major biological groups. For example, Na could be predicted with *R^2^* = 0.519, but mainly because Australian *A. flexuosa* has on average more than twice the Na concentration of other species sampled. The model could have incorporated any optical features that distinguish *A. flexuosa* from other specimens, whether or not those features truly result from Na concentration. This kind of constellation effect may underlie the spectroscopic estimation of many elements (Nunes et al. 2017; Kothari & Schweiger 2022).

### Interpreting PLSR models

The VIP metric revealed strong similarities across traits in the major peaks of importance, including the green hump, the red edge, and multiple features in the SWIR (e.g. 1390 and 1880 nm; Fig. S13-15). The general pattern of high VIP along the green hump and at the red edge is shared by many traits across studies (Yang et al. 2016; Ely et al. 2019; Yan et al. 2021; Kothari et al. 2022). In general, visible and red edge reflectance are dominated by pigments and leaf structure (Richardson et al. 2002). Many traits show isolated VIP peaks in the SWIR, which are harder to explain in terms of specific traits. Two peaks are in off-center parts (1390 nm and 1880 nm) of major water absorption features, and the SWIR also contains many shallow, overlapping absorption features of bonds in dry matter components, making it hard to isolate the influence of any single one (Curran et al. 1989). In contrast to these distinct peaks, LMA and EWT (as well as some elements) had high VIP throughout much of the SWIR, which might arise from the strong and pervasive influence of LMA and EWT on SWIR reflectance.

Similarities in VIP may result from trait covariance (Fig. S1), which could allow models to leverage constellation effects. Some cases are clear: the three pigment pools correlate tightly with each other (*R^2^* = 0.853-0.974) and with N (*R^2^* = 0.457-0.517), and their VIP patterns share an overall visual similarity. Likewise, solubles, hemicellulose, and cellulose are closely linked (*R^2^* = 0.643-0.834) and have similar VIP patterns. Other explanations are more elusive: for example, all four carbon fractions have high VIP across the green hump and red edge, but correlate poorly (*R^2^* = 0-0.107) with pigments and LMA, which account for most spectral variation in those regions. Such findings imply that strong covariance with a few key traits cannot alone explain the striking similarities in VIP, nor the high accuracy of models for traits like carbon fractions in the internal validation.

### Alternate PLSR models

We focused on estimating chemical traits on a mass basis, but the remote sensing literature contains powerful arguments for estimating certain constituents on an area basis. In particular, the optical properties of leaves are driven more by the absolute quantity of a constituent than its quantity relative to other constituents. This consideration should matter most for traits distributed in proportion to area (Kattenborn et al. 2019). In our dataset, pigments and several nutrients were mostly area-proportional (Table S1). As Kattenborn et al. (2019) suggested, estimating area-based traits directly was usually more accurate than multiplying mass-based estimates by LMA estimates, especially for traits distributed in proportion to area (Table S1). This consideration is important given that many research questions may require traits expressed on an area basis.

Using transmittance or absorptance spectra instead of reflectance spectra increases accuracy for most traits, but only slightly (Table S2). It may be relevant that the traits that show the greatest improvement include most of the highly area-proportional traits. Given that transmittance or absorptance spectra yielded little improvement for most traits, adding transmittance measurements may not aid much in trait estimation, although they are still useful in endeavors like radiative transfer modeling (Spafford et al. 2021).

### External validation and model transferability

Aside from our internal validation, we also considered model performance under less ideal scenarios: we tested the primary models on external validation datasets, and built another set of models to be tested on a functional group left out during their construction. When transferring models to left-out functional groups, performance was usually worse (lower *R^2^*, higher RMSE_full_) than when models were evaluated on a random internal validation dataset (Fig. 6; Table S5). This finding is consistent with previous studies transferring models among species (Helsen et al. 2021) or sites (Yan et al. 2021). In general, trait models performed worse when the trait distributions in the calibration and validation datasets were less similar, but the strength and even the direction of the trend varied among traits (Fig. 6B). Thus, support for Hypothesis 1 was equivocal. Predictions of LMA among conifers and carbon fractions among graminoids were strongly biased and failed to capture the functional distinctiveness of these groups. These results underscore that models must be built using samples whose diversity encompasses the models’ expected domain of application (Yang et al. 2016; Schweiger 2020; Beauchamp-Rioux 2022).

The external validation was a more severe test of model generality because the data were collected on different species with different instruments using different trait measurement protocols. This kind of scenario may better represent a realistic use case for such models. All traits showed declines in performance from internal to external validation (Table 2). This result is expected and consistent with previous findings: for example, Serbin et al. (2019)’s global model of LMA showed *R^2^* = 0.89 for internal validation and 0.66 for external validation on LOPEX and ANGERS, as compared to *R^2^* = 0.89 (internal) and 0.59 (external, averaged across datasets) here. These results further underscore the challenges that arise when transferring models among settings and instruments.

The rank-order of *R^2^* across traits was broadly similar between the internal validation, external validation, and model transferability analyses (Fig. 6A), which offers some general support for Hypothesis 2. Models for LMA, LDMC, and EWT performed relatively well in all analyses, while models for most macro- and micronutrients performed poorly in both internal and external validation (but were not included in model transferability analyses). Pigments, N, C, and carbon fractions were generally intermediate. However, in both the external validation and model transferability analyses, C and carbon fractions showed particularly strong declines in *R^2^* compared to internal validation (Fig. 6A). Dry matter components like cellulose and lignin are associated mainly with weak absorption features throughout the NIR and SWIR (Curran et al. 1989). In contrast, traits without such strong declines—LMA, LDMC, EWT, N, and pigments—have stronger influences on the spectrum, and many are part of mechanistic models like PROSPECT (Féret et al. 2008). This contrast could imply that models are most transferable when their target traits have strong absorption features. Many chemical traits may be estimated more reliably using spectra of pressed or ground leaves, which by removing the water absorption features expose the subtler features of dry matter components (Kothari et al. 2022).

Even though our estimates of LMA and EWT correlated well with measured values in the external validation, they also had a strong bias towards overestimation (Fig. 5), which could complicate their use in practice. This bias may have resulted from varying measurement protocols, or from differences in instrumentation that can have a subtle but pervasive influence on the measured spectra (Castro-Esau et al. 2006; Lukeš et al. 2017; Hovi et al. 2018). This bias could perhaps be corrected by measuring a subset of trait values to determine how they correspond to model predictions, and then transforming the remaining predictions accordingly. In the longer term, it might become easier to get accurate trait estimates using existing models if techniques are developed to reconcile differences among instruments (Schweiger 2020; Meireles et al. 2020). Deep learning approaches like convolutional neural networks may also address this problem through their greater flexibility and capacity for transfer learning (Kattenborn et al. 2021).

### Implications for global trait-spectra modeling

Our models draw from samples representing several functional groups and a wide range of temperate ecosystems, including wetlands, grasslands, and closed-canopy forests (Table 1). However, these samples represent just a portion of Earth’s vast plant diversity—omitting, for example, tundra, drylands, and tropical biomes. These biomes would surely contain combinations of optically important traits missing in our dataset, which would need to be represented in a universal model that could be used in mapping functional trait variation worldwide. We hope plant ecologists can use our data and models not only as a tool to address ecological questions, but also as a foundation for building yet more general models.

Our results imply that some traits can consistently be estimated better than others. We can compare our findings to Asner et al. (2011), who measured a similar set of traits on a vast number of samples from canopy leaves in humid tropical forests. The two studies are broadly consistent despite representing disparate subsets of the world’s plant diversity. Both found that models for LMA and water content were most accurate (*R^2^*> 0.85), followed by, in varying orders, C and N, pigments, and carbon fractions (*R^2^*= 0.55-0.85). In both studies, models for other elements were typically worse (*R^2^*< 0.65), but better for Ca and K than many other elements. Both studies also found that transmittance spectra yielded slightly better performance for most traits. The stability in rankings suggests a general hierarchy of traits that are the most practical targets for reliable model-based estimation.

The emerging body of research showing that spectral models can yield fast, reliable estimates of many leaf traits across functional groups and ecosystems could expand our ability to map and monitor plant function around the world, bridging between conventional trait measurements and remote sensing. Our findings clarify both the opportunities and challenges of using spectral trait estimates. Research has not yet progressed to the point where plant ecologists can use existing empirical models to estimate traits in their study systems without further validation. Aside from the issues of taxonomic and functional representativeness, our external validation underscores the challenges of transferring models among instruments, which might require new methodological developments to overcome. Even under ideal conditions, some aspects of leaf function (e.g. elemental composition) may be hard to measure accurately based on spectroscopy alone. We hope that the promise of rapid, large-scale monitoring of plant function inspires the coordinated effort needed to understand and (when possible) overcome these challenges.

In sum, we built models to predict traits from reflectance spectra using nearly 2000 leaf samples from 104 species. Traits varied a great deal how well they could be estimated using spectral models. Our models for structural and water-related traits were very accurate in internal validation and showed great promise in external validation; in contrast, our models for many micronutrients performed very poorly. We suggest that these patterns reflect a general hierarchy that emerges mainly from the varying degrees of influence these traits have on the spectrum. Leaf reflectance spectra thus bear the imprint of many aspects of leaf function—but not all to the same degree. Most importantly, we show that many traits can be estimated accurately from spectra using general models that span multiple functional groups and ecosystems. This finding represents a significant advance in leveraging spectra towards major goals in functional ecology.

## Supporting information

All supplementary material

## Acknowledgements

We conducted this research at institutions and field sites located on the largely unceded ancestral and contemporary land of many First Nations and Aboriginal people. We thank Jocelyne Ayotte, Zachary Bélisle, Fabien Cichonski, Chris Colton, Myriam Cloutier, Aurélie Dessain, Vincent Fournier, Juliette Frappier-Lecomte, Isabelle Gareau, Sam Grubinger, Elisabeth Hardy-Lachance, Sandra Jooty, Alexandra Massey, Antoine Mathieu, Chris Mulverhill, Rime Néron, Florence Normand-Boisseau, Clement Robert-Bigras, Sabine St-Jean, Guillame Tougas, Charlotte Taillefer, Francois du Toit, and Madeleine Trickey-Massé for their help with field sampling and laboratory analyses. We thank the National Capital Commission, the Société des établissements de plein air du Québec, the Station de biologie des Laurentides of Université de Montréal, the Jardin botanique de Montréal, the Institut de recherche d’Hydro-Québec, the Nature Conservancy of Canada, and the Parks and Wildlife Service of Western Australia for permissions to conduct sampling and for lending the ecological and logistical expertise of their staff. We also thank the many members of CABO have helped to strengthen this work through their stimulating comments and questions. CABO was funded by a Discovery Frontiers grant to EL, MK, AB, MV and NCC from the Natural Sciences and Engineering Research Council of Canada (NSERC; grant number 509190-2017).

## Author contributions

SK and EL designed the project with contributions from AB, NCC, MK, and MV. RBR, FB, ALC, AG, XGM, PWH, JP, AKS, and SDT collected and curated the spectral and trait data. SK analyzed the data, interpreted the results, and wrote the first draft with substantial contributions from EL. All authors contributed to further revisions of the paper.

## Data availability

All fresh-leaf spectral data are available through the CABO data portal (https://data.caboscience.org/leaf). Upon publication, we will also upload all spectral data, as well as metadata and trait data, to the Ecological Spectral Information System (EcoSIS, https://ecosis.org/), and upload models to the Ecological Spectral Model Library (EcoSML, https://ecosml.org/). At that stage, we will update this section accordingly. Analysis scripts are available as a repository on GitHub (https://github.com/ShanKothari/CABO-trait-models).

