## Supplementary material for "Predicting leaf traits across functional groups using reflectance spectroscopy": All supplementary material

**Supplementary Materials**

*Sampling*

We collected spectra and traits for all specimens using consistent protocols. For each sample, we collected sunlit leaves (>3 h per day of sun exposure) that were relatively homogeneous—for trees and shrubs, at about the same canopy position in the same individual. When possible, we aimed to collect enough leaves in each sample to carry out all destructive trait measurements while retaining several grams of leaf tissue in long-term storage. We avoided leaves that showed clear signs of senescence, and (particularly for spectral measurements) aimed to choose healthy-looking leaves without visible herbivore or pathogen damage. On a subset of leaves, we measured reflectance and transmittance spectra and punched leaf disks for pigment analyses. We also selected random subsets of visually similar leaves to measure other structural or chemical traits. To prepare each sample for chemical trait measurements, we dried several grams of leaves at 65 °C for three days and ground them in a 2 mm cyclone mill. All trait measurements exclude petioles but include raches of compound leaves, since they are functionally equivalent to the midrib of a simple leaf.

We identified species in the field and used the Database of Vascular Plants of Canada (VASCAN; Desmet & Brouillet 2013) to classify each species by its family and growth form. VASCAN classified plants in our dataset as either shrubs, trees, herbs, or vines. Some species were listed as adopting either shrub or tree growth forms, in which case we manually classified them based on the habit of the individuals sampled. We further divided species into functional groups as follows: we classified trees as either needleleaf coniferous trees (here, families Pinaceae or Cupressaceae) or broadleaf trees; and we classified herbs as either graminoids (families Poaceae, Cyperaceae, Typhaceae, or Juncaceae) or forbs. The Australian species *Agonis flexuosa* (Willd.) Sweet was not in VASCAN, so we assigned it to the broadleaf tree functional group. Broadleaf trees were by far the best-represented functional group ($n$ = 996; species = 42), but the database also includes graminoids ($n$ = 359; species = 15), forbs ($n$ = 242;

species = 20), shrubs (*n* = 223; species = 20), conifers (*n* = 128; species = 12), vines (*n* = 16; species = 1), and ferns (*n* = 7; species = 1). In total, the database includes samples from 31 families.

*Spectral measurements and processing*

We measured directional-hemispherical reflectance and transmittance spectra (350-2500 nm) using an HR-1024i spectroradiometer equipped with a DC-R/T integrating sphere from Spectra Vista Corporation (Poughkeepsie, NY, USA). The instrument's nominal bandwidth varies among sensors: ≤ 1.5 nm through the range of the silicon detector array (~350-1000 nm), ≤ 3.8 nm through the first InGaAs array (~1000-1890 nm), and ≤ 2.5 nm through the second InGaAs array (~1890-2500 nm). The spectral resolution likewise ranges from 3.3 to 9.5 nm, depending on the spectral region. For each sample, we measured spectra from the adaxial surface of six leaves or leaf arrays (Laliberté & Soffer 2018a). For leaves that were too small or narrow to cover the port of the integrating sphere, we followed a protocol adjusted from Noda et al. (2013) to measure single-layer composites of leaves (Laliberté & Soffer 2018b). We either sealed leaves in plastic bags or left them attached to the branch to reduce water loss during the short interval between collection and measurement. Reflectance measurements were calibrated against a white Spectralon 99% reflectance standard and corrected for stray light.

We processed reflectance and transmittance spectra following Schweiger & Laliberté (2020). We resampled the spectra linearly to 1 nm resolution and interpolated linearly over the sensor overlap regions. We then averaged spectra from leaves of the same sample and reduced noise by applying a Savitzky-Golay filter with varying order and length: order 3 and length 21 from 350-715 nm, order 3 and length 35 from 715-1390 nm, order 3 and length 75 from 1390-1880 nm, and order 5 and length 175 from 1880-2500 nm. We trimmed spectra to 400-2400 nm to remove the particularly noisy ends of the spectrum. Finally, we calculated absorptance at each wavelength by subtracting the sum of reflectance and transmittance from 1. We performed all spectral processing using *spectrolab* v. 0.0.10 (Meireles et al. 2017) in *R* version 3.6.3 (R Core Team 2020).

Spectral measurement protocols are given in other sources for external validation data, including LOPEX (Hosgood et al. 1993; Hosgood et al. 1994), ANGERS (Jacquemoud et al. 2003; Féret et al. 2008; http://opticleaf.ipgp.fr/index.php?page=database), and Dessain (Kothari et al. 2022). The Dessain dataset is expanded from our previous description in Kothari et al. (2022); it has 200 samples, including trees ($n$ = 107; species = 36), shrubs ($n$ = 53; species = 29), forbs ($n$ = 28; species = 19), graminoids ($n$ = 6; species = 2), ferns ($n$ = 4; species = 3), and vines ($n$ = 2; species = 2). Each dataset contained spectra that included the 400-2400 nm range already resampled to 1 nm resolution. For consistency, we processed the external validation spectra much like our core dataset. First, we either interpolated over the sensor overlap region (Dessain) or matched sensors using *match_sensors()* in *spectrolab*. Based on a visual assessment, we matched sensors at 862 nm for LOPEX, but did not match sensors for ANGERS because there was no conspicuous offset at the sensor overlap region. Next, we applied a Savitzky-Golay filter to the three datasets, using the same parameters we used on the CABO dataset. Finally, we trimmed all spectra to 400-2400 nm.

*Trait measurements*

Leaf mass and area

We measured the fresh mass of a subset of leaves from each sample in the field shortly after collection. We rehydrated the leaves in sealed plastic bags with damp paper towels and stored them in a refrigerator for at least 12 h. We then measured their rehydrated mass, scanned them, and measured their area using WinFOLIA (Regent Instruments, Québec, QC, CA). Finally, we dried them at 65 °C for at least 72 h before measuring their dry mass. We used total masses and areas, summed across all leaves from a sample, in trait calculations. We calculated leaf mass per area (LMA) as the dry mass divided by leaf area; leaf dry matter content (LDMC) as dry mass divided by fresh mass; and equivalent water thickness (EWT) as fresh mass minus dry mass, all divided by leaf area (Laliberté 2018).

Chemical traits

We estimated leaf carbon and nitrogen concentration on ground samples using an Elementar Vario MICRO Cube (Langenselbold, Hesse, Germany; Ayotte et al. 2019). We measured concentrations of ten other nutrients (Al, Ca, Cu, Fe, K, Mg, Mn, Na, P, and Zn) using nitric acid digestion followed by inductively coupled plasma optical-emission spectrometry (ICP-OES) on an Optima 7300DV (PerkinElmer, Waltham, MA, USA). These ICP-OES data are available only for the Beauchamp-Rioux, Boucherville, Girard, and Warren projects. We removed six data points from the ICP-OES output as unrepresentative outliers because their reported values were >30% greater than all others for their respective elements. We also set concentrations below the detection limit, which sometimes returned as negative, to zero.

We measured carbon fractions on ground samples using an ANKOM2000 Fiber Analyzer (ANKOM Technology, Macedon, NY, USA; Ayotte & Laliberté 2019). The procedure is a sequential digestion in a series of solutions that wash away fractions of the leaf tissue, leaving behind fractions that are increasingly recalcitrant. The first digestion at 100 °C in a neutral detergent solution washes away soluble cell contents. The second digestion at 100 °C in a mildly acidic detergent solution washes away bound proteins and hemicellulose, which we simply call hemicellulose. The third digestion at room temperature in 72% sulfuric acid removes cellulose, leaving behind lignin and other recalcitrant materials. Finally, we ashed all samples in a furnace to separate inorganic recalcitrant materials (ash) from organic ones, which we simply call lignin. By weighing the remaining leaf tissue after each stage, we could determine the mass of each fraction. We excluded solubles and hemicellulose from three species (*Acer platanoides* Linnaeus, *Ulmus americana* Linnaeus, and *Ulmus rubra* Muhlenberg) and all carbon fractions from *Alnus incana subsp. rugosa* (Du Roi) R.T. Clausen because of leaf properties that caused unreliable results. For example, *A. platanoides* has leaf exudates that wash off inconsistently in neutral detergent; *A. incana* samples often returned negative values of hemicellulose.

We stored leaf disks in a cooler with ice in the field and transferred them first to a -20 °C freezer in the lab, then to a -80 °C freezer for long-term storage. When we needed to ship them (e.g. the Warren project), we freeze-dried them first. We extracted pigments from frozen leaf disks (or cut pieces, for narrow-leaved plants) under darkness in an $MgCO_3$-MeOH solution and estimated pigment concentrations on a SPECTROstar Nano microplate reader (BMG LABTECH, Ortenburg, Germany) using a spectrophotometric protocol (Girard et al. 2020). While concentrations were measured per unit fresh mass, we used LDMC as a conversion factor to express them per unit dry mass.

We occasionally removed specific trait values when an error was noted during the measurement protocol.

External validation

Trait measurement protocols are given in the aforementioned sources for external validation data, including LOPEX (Hosgood et al. 1994), ANGERS (Féret et al. 2008; http://opticleaf.ipgp.fr/index.php?page=database), and Dessain (Kothari et al. 2022). In the Dessain dataset, we did not measure initial fresh mass of leaves and instead used the rehydrated mass. This difference could cause EWT to be overestimated and LDMC to be underestimated. However, relative water content in the remaining projects was usually close to 100% (median: 87.7%, 2.5–97.5$^{th}$ percentile: 68.3–97.6%). We also removed solubles and hemicellulose values from the Dessain dataset for the same species whose values we removed from the main CABO dataset. Finally, we used a conservative outlier detection procedure—the same as we used for the main CABO dataset—to remove eight data points from Dessain ICP-OES data. In LOPEX, there were two estimates of both cellulose and lignin for each sample, so we took the average for each one.

Besides applying the main set of models described in the main text, we also tested whether certain transformations of the spectral data could improve performance on the external validation data—as, for example, if certain instruments tended to measure higher or lower reflectance across the spectrum. We used: (1) brightness normalization, which normalizes each spectrum to a unit vector while preserving its shape (Feilhauer et al. 2010), and (2) continuum removal, which performs an albedo normalization by reporting the difference between the actual spectrum and a linear interpolation of its convex hull (Clark & Roush 1984). We first transformed the spectra in the CABO dataset using either brightness normalization or continuum removal and built models using the calibration and validation approaches described in the main text (model performance for internal validation not shown). We then applied the models to the external validation data transformed likewise. Neither transformation caused any systematic improvement in trait estimates (Tables S3-4).

**Supplementary figures**

**Fig. S1:** (top) The first two dimensions in a principal components analysis (PCA) of CABO trait data, excluding all elements other than C and N. All chemical trait data are expressed on a mass basis, as in the core PLSR analyses of this paper. (bottom) A PCA of CABO trait data with chemical trait data expressed on a normalization-independent basis (Osnas et al. 2013), which is meant to eliminate covariance induced by jointly normalizing variables by mass. Here, we also remove solubles, since carbon fractions are nearly constrained to sum to 1, as well as chlorophyll *b* and carotenoids, since the three pigment pools covary strongly.

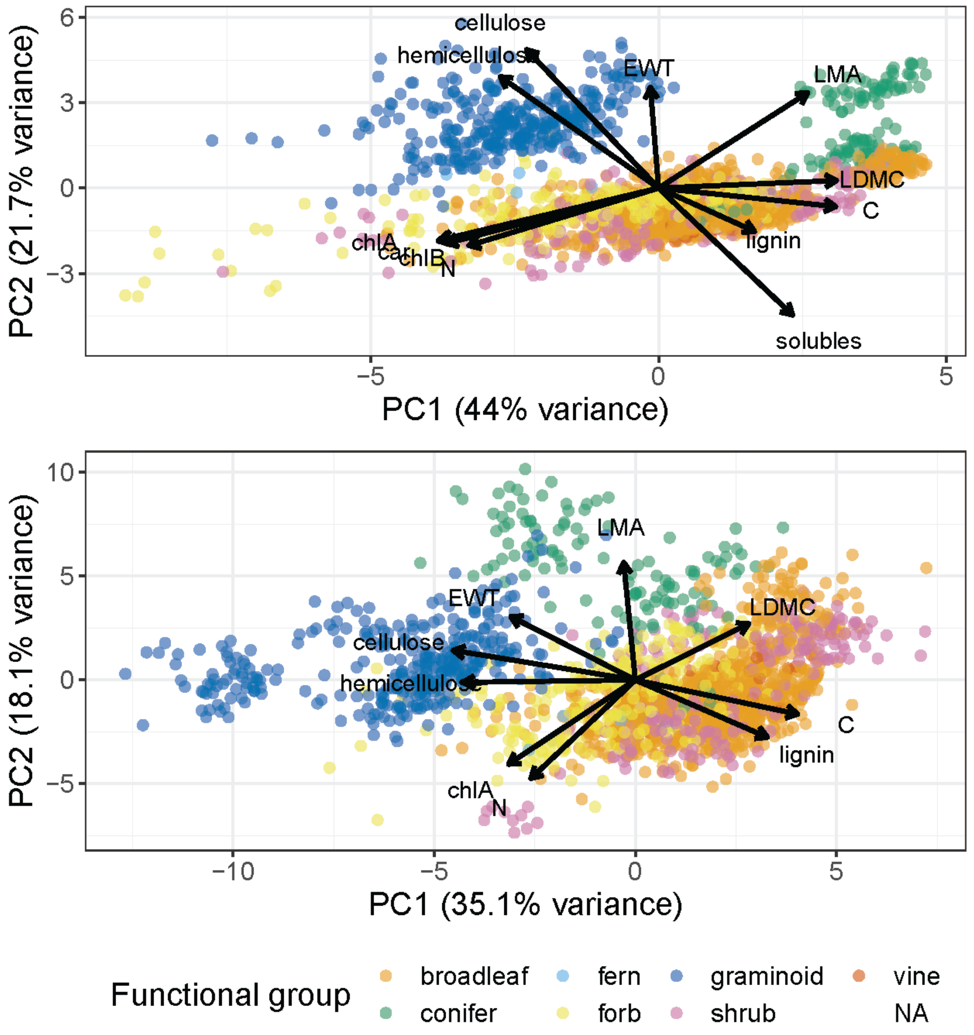

**Fig. S2:** The median leaf-level (A) reflectance, (B) transmittance, and (C) absorptance at each wavelength across the spectrum, separated by functional group. We omit ferns and vines, which are only represented by one species each. The dashed line shows the coefficient of variation across all samples.

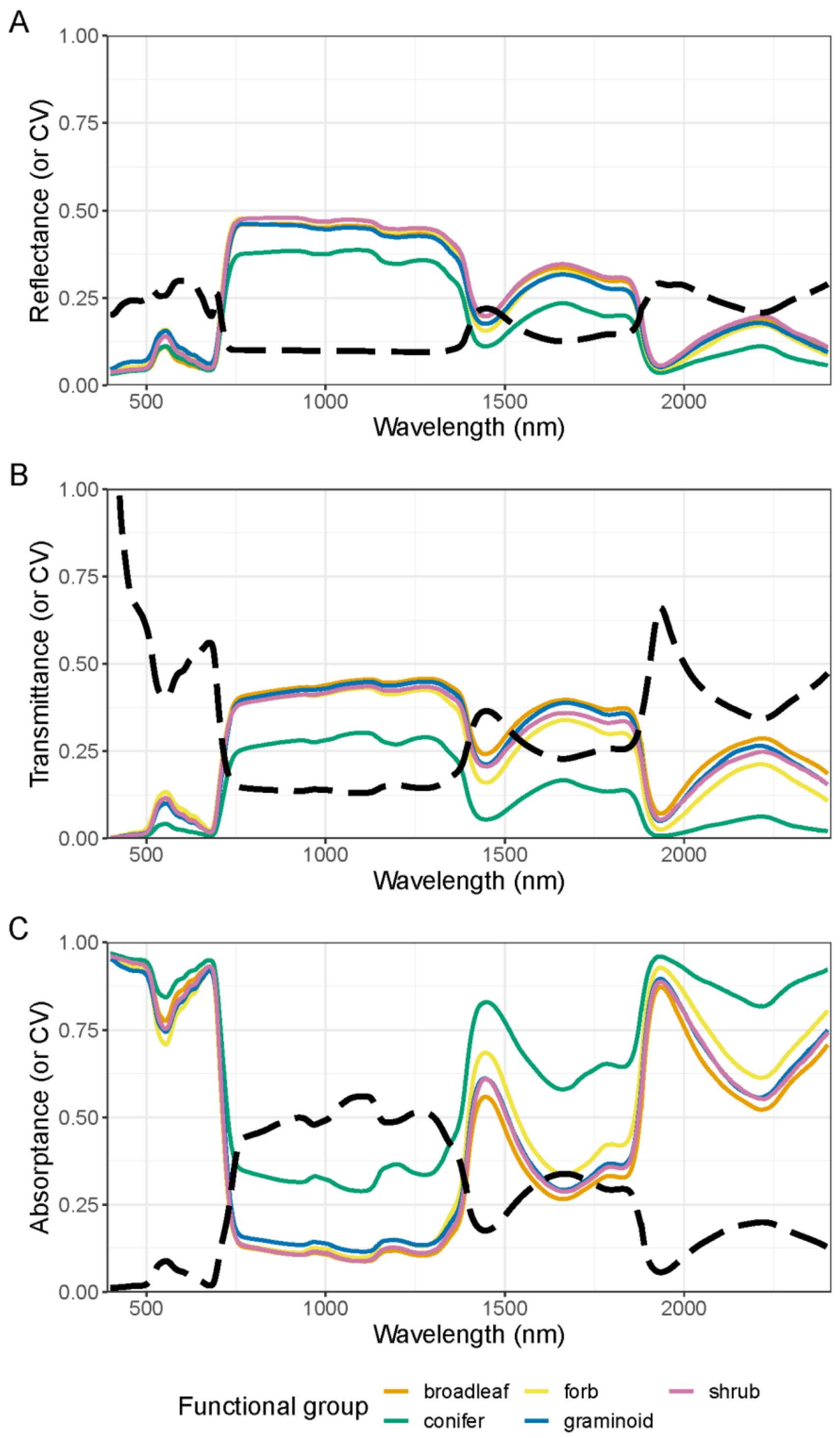

**Fig. S3:** Plots of observations against PLSR predictions among internal validation data for structural and water-related traits, comparing predictions from reflectance, transmittance, and absorptance spectra. The black dashed line in each panel is the 1:1 line. Colored lines represent best-fit lines from OLS regression for each functional group. Error bars around each point represent 95% confidence intervals based on the ensemble of models produced in the 100× jackknife analysis.

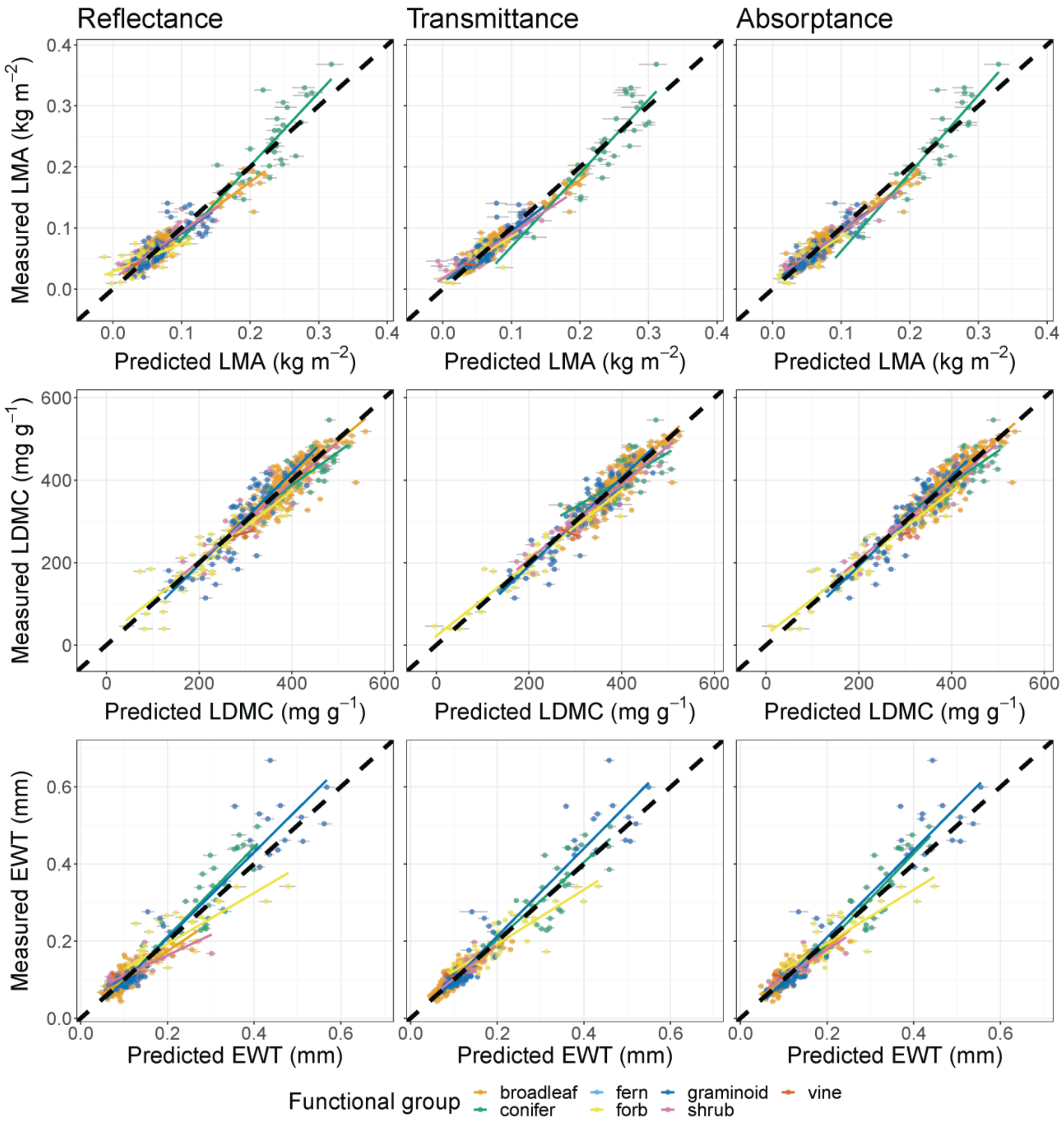

**Fig. S4:** Plots of observations against PLSR predictions among internal validation data for N, C, and P, comparing predictions from reflectance, transmittance, and absorptance spectra. P is only available for the Beauchamp-Rioux, Boucherville 2018, Girard, and Warren projects. The black dashed line in each panel is the 1:1 line. Colored lines represent best-fit lines from OLS regression for each functional group. Error bars around each point represent 95% confidence intervals based on the ensemble of models produced in the 100× jackknife analysis.

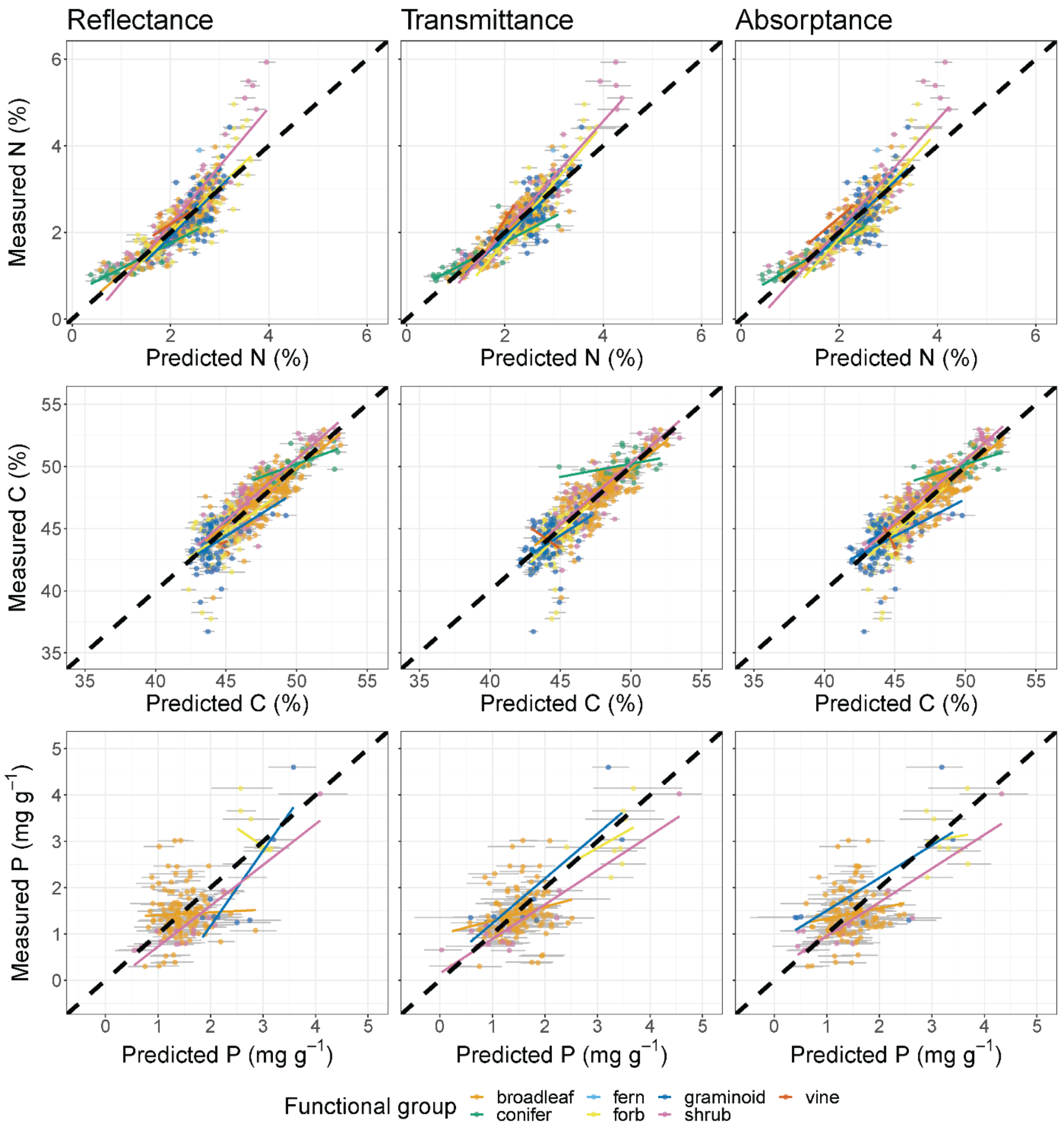

**Fig. S5:** Plots of observations against PLSR predictions among internal validation data for carbon fractions, comparing predictions from reflectance, transmittance, and absorptance spectra. The black dashed line in each panel is the 1:1 line. Colored lines represent best-fit lines from OLS regression for each functional group. Error bars around each point represent 95% confidence intervals based on the ensemble of models produced in the 100× jackknife analysis.

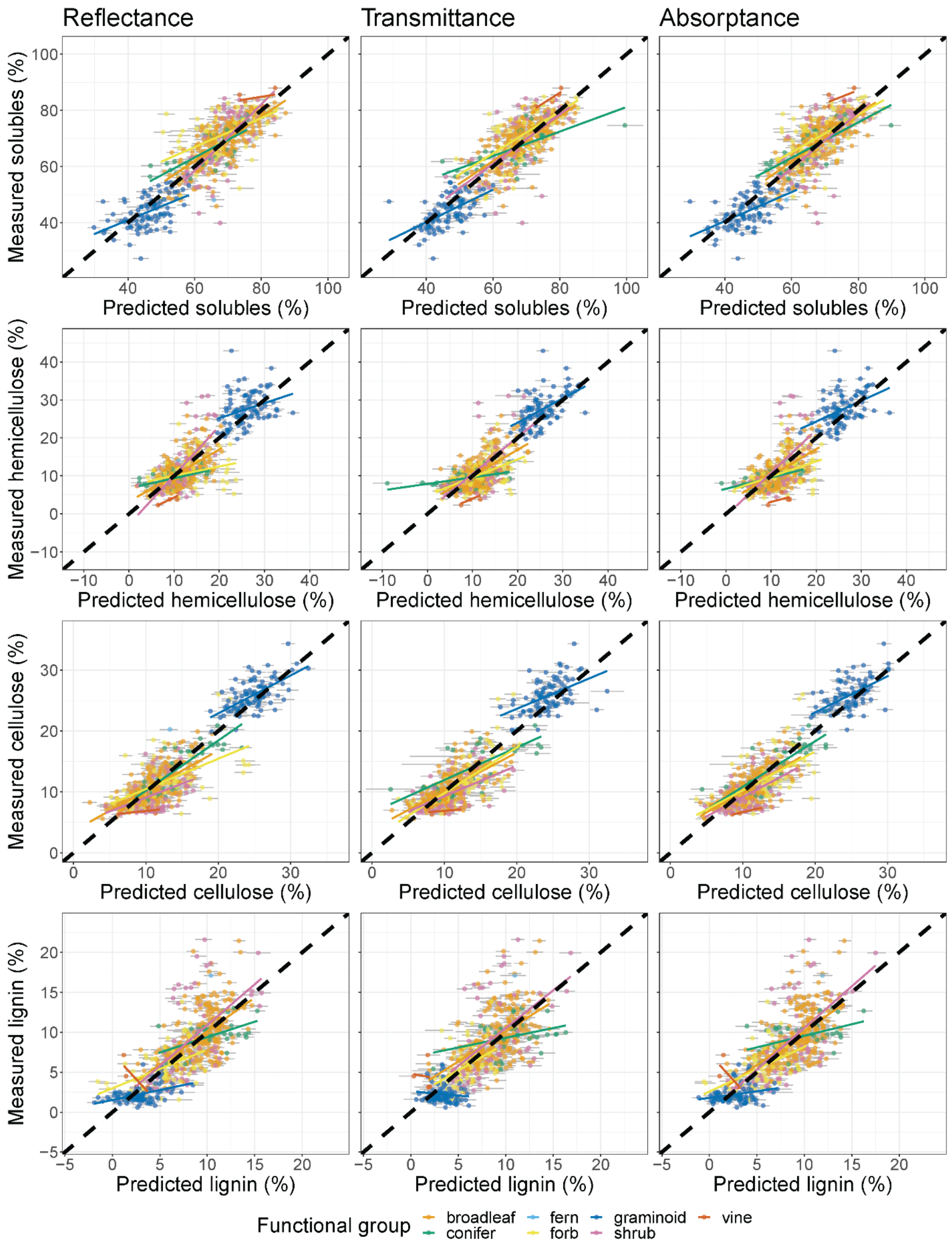

Reflectance
Transmittance
Absorptance
Measured solubles (%)
Predicted solubles (%)
Measured hemicellulose (%)
Predicted hemicellulose (%)
Measured cellulose (%)
Predicted cellulose (%)
Measured lignin (%)
Predicted lignin (%)
Functional group
broadleaf
conifer
fern
forb
graminoid
shrub
vine

**Fig. S6:** Plots of observations against PLSR predictions among internal validation data for pigments, comparing predictions from reflectance, transmittance, and absorptance spectra. The black dashed line in each panel is the 1:1 line. Colored lines represent best-fit lines from OLS regression for each functional group. Error bars around each point represent 95% confidence intervals based on the ensemble of models produced in the 100× jackknife analysis.

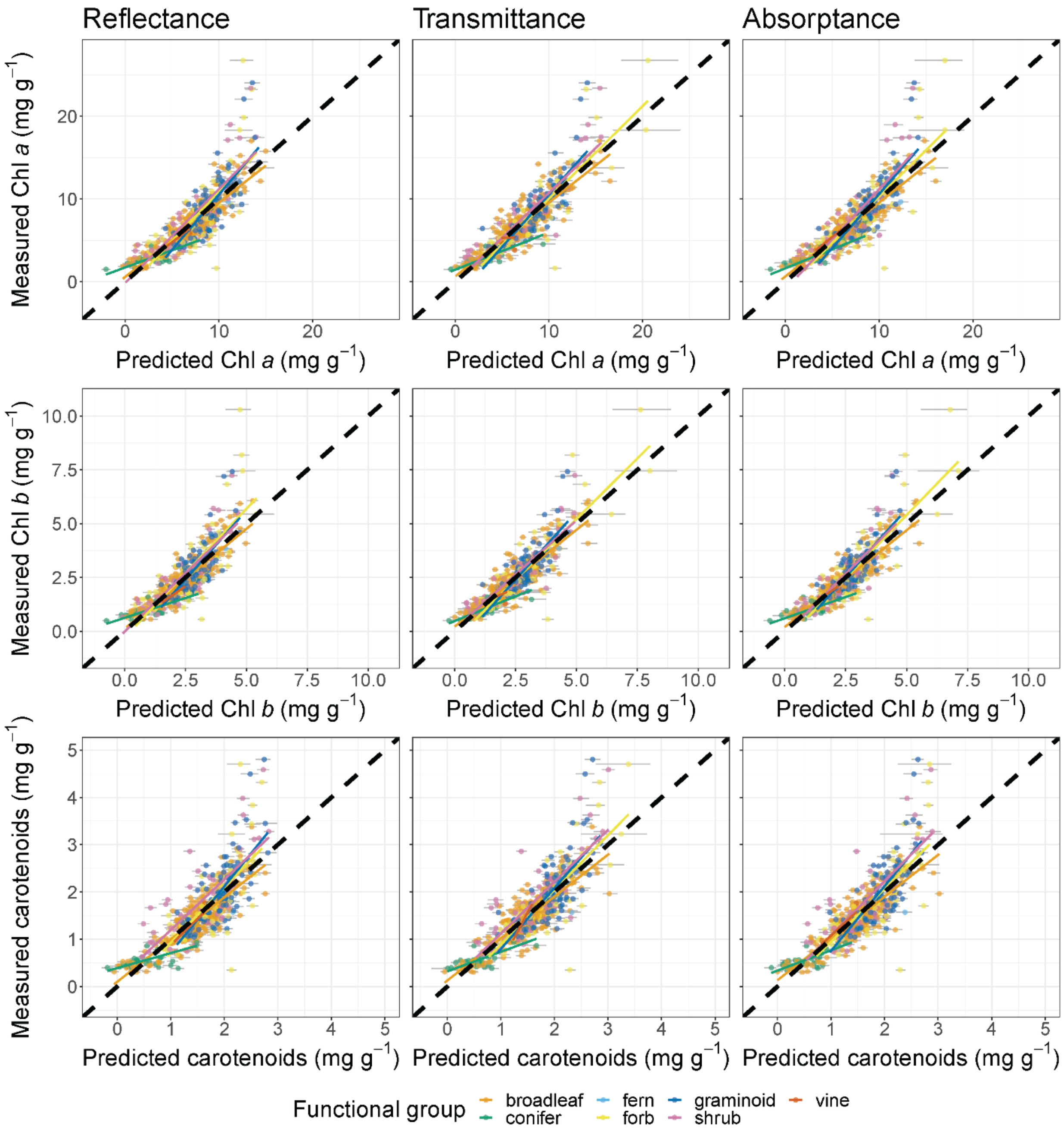

**Fig. S7:** Plots of observations against PLSR predictions among internal validation data for Al, Ca, and Cu, comparing predictions from reflectance, transmittance, and absorptance spectra. These data are only available for the Beauchamp-Rioux, Boucherville 2018, Girard, and Warren projects. The black dashed line in each panel is the 1:1 line. Colored lines represent best-fit lines from OLS regression for each functional group. Error bars around each point represent 95% confidence intervals based on the ensemble of models produced in the 100× jackknife analysis.

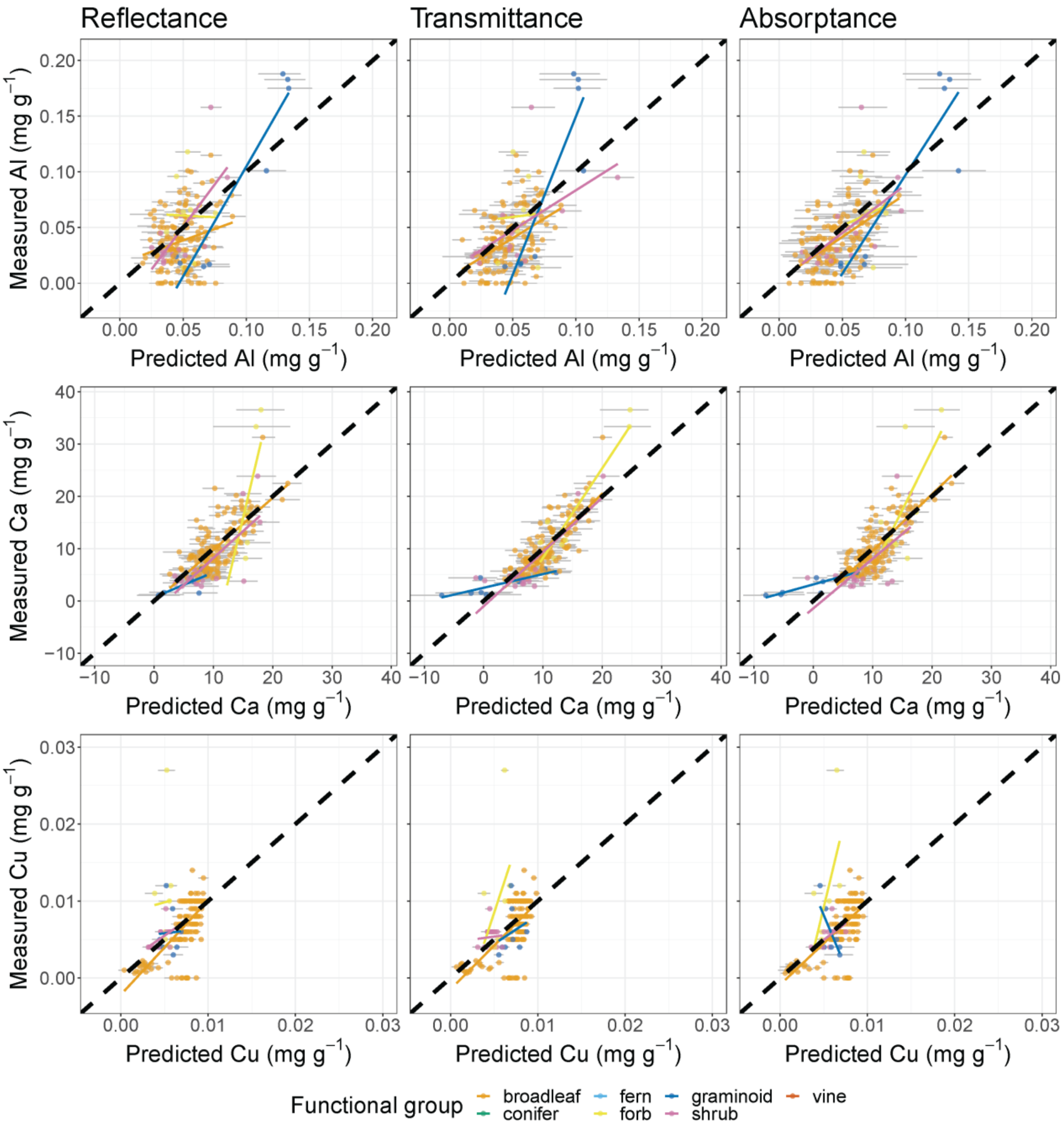

**Fig. S8:** Plots of observations against PLSR predictions among internal validation data for Fe, K, and Mg, comparing predictions from reflectance, transmittance, and absorptance spectra. These data are only available for the Beauchamp-Rioux, Boucherville 2018, Girard, and Warren projects. The black dashed line in each panel is the 1:1 line. Colored lines represent best-fit lines from OLS regression for each functional group. Error bars around each point represent 95% confidence intervals based on the ensemble of models produced in the 100× jackknife analysis.

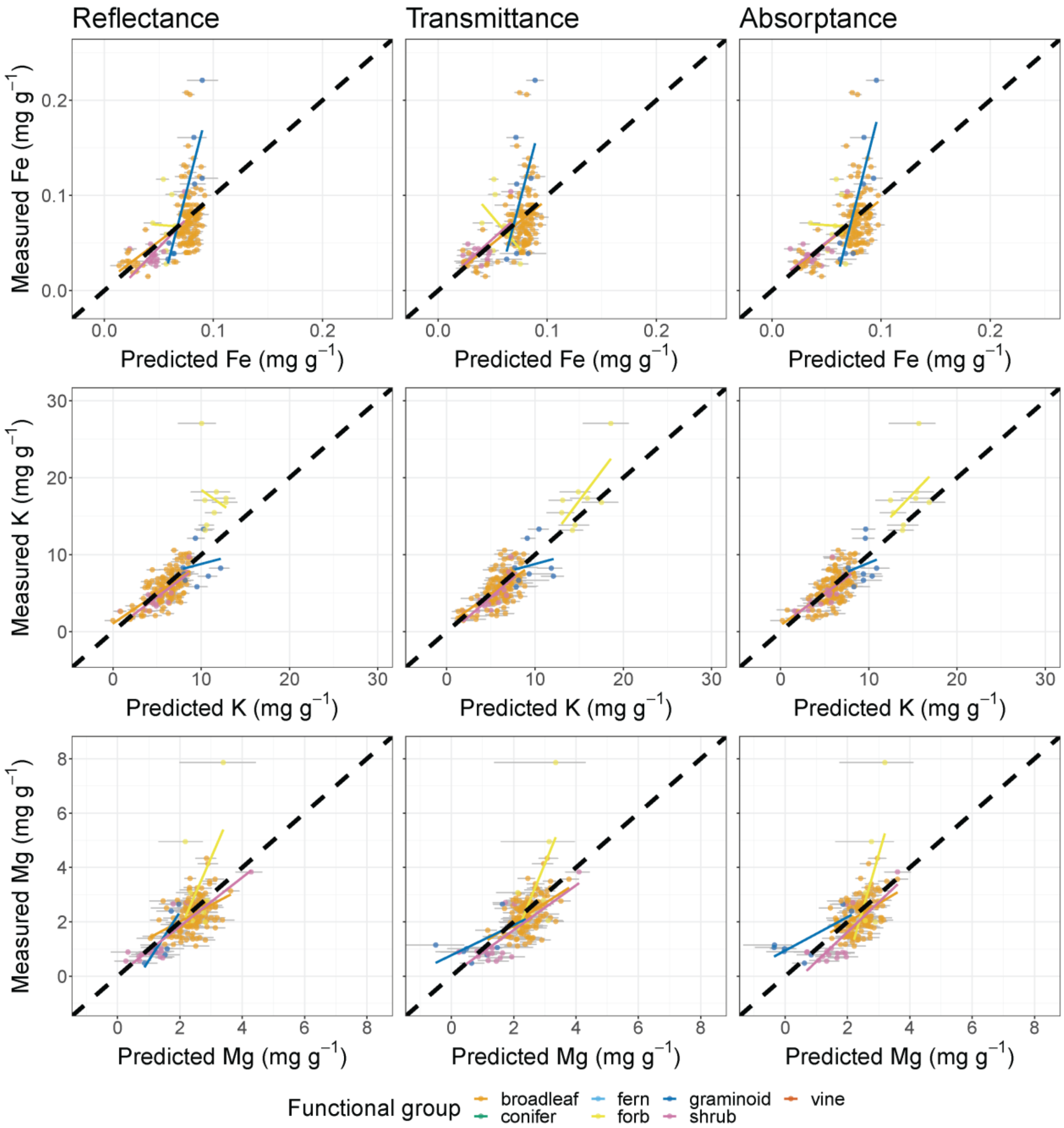

**Fig. S9:** Plots of observations against PLSR predictions among internal validation data for Mn, Na, and Zn, comparing predictions from reflectance, transmittance, and absorptance spectra. These data are only available for the Beauchamp-Rioux, Boucherville 2018, Girard, and Warren projects. The black dashed line in each panel is the 1:1 line. Colored lines represent best-fit lines from OLS regression for each functional group. Error bars around each point represent 95% confidence intervals based on the ensemble of models produced in the 100× jackknife analysis.

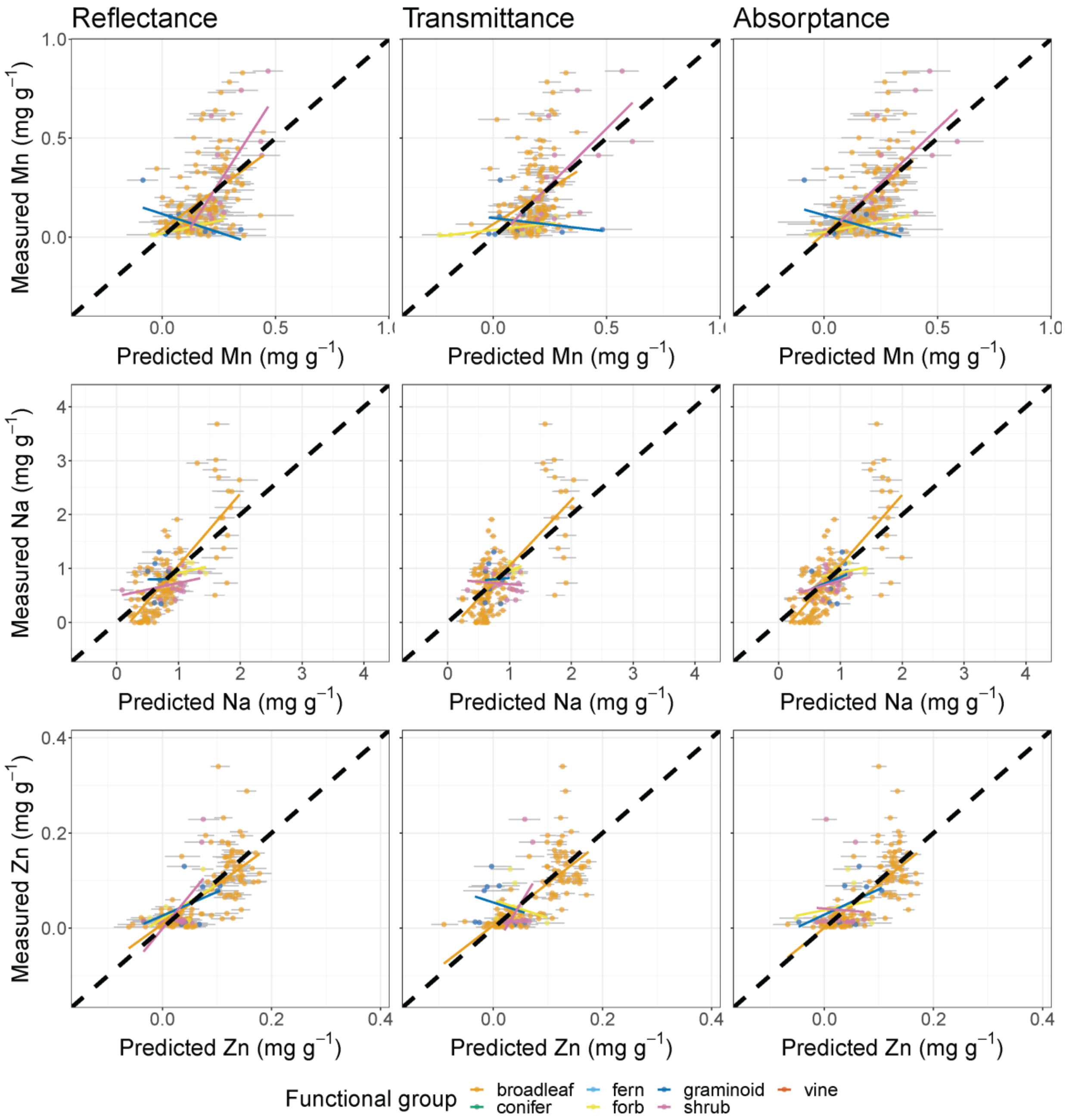

**Fig. S10:** The distributions of reflectance-based model performance statistics for each trait based on 100 jackknife resampling iterations from the calibration data set. The red dots show the $R^2$ and %RMSE from applying the ensemble of models to the internal validation subset. Abbreviations: sol = solubles, hemi = hemicellulose, cell = cellulose, lign = lignin, car = total carotenoids.

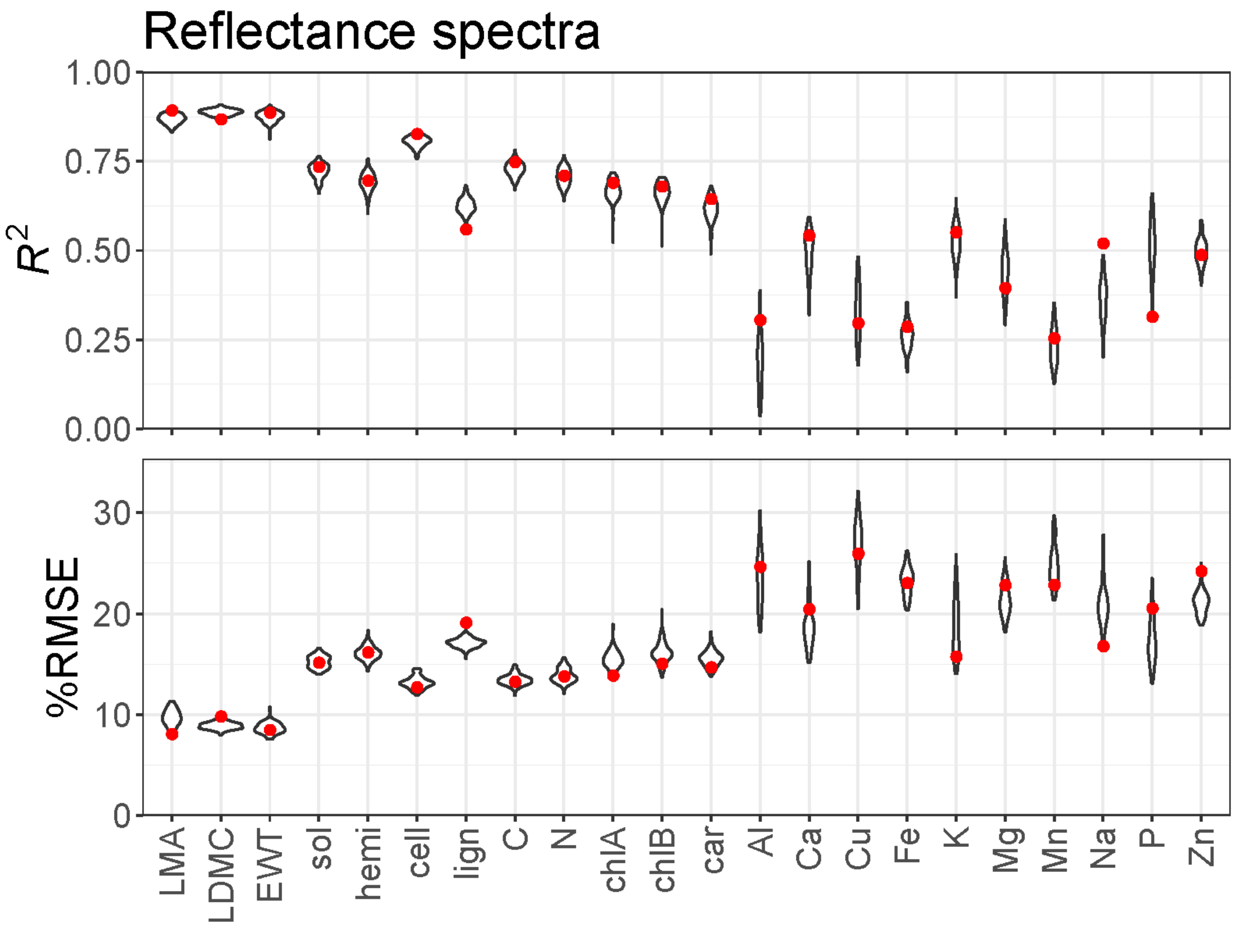

**Fig. S11:** The distributions of transmittance-based model performance statistics for each trait based on 100 jackknife resampling iterations from the calibration data set. The red dots show the $R^2$ and %RMSE from applying the ensemble of models to the internal validation subset. Abbreviations: sol = solubles, hemi = hemicellulose, cell = cellulose, lign = lignin, car = total carotenoids.

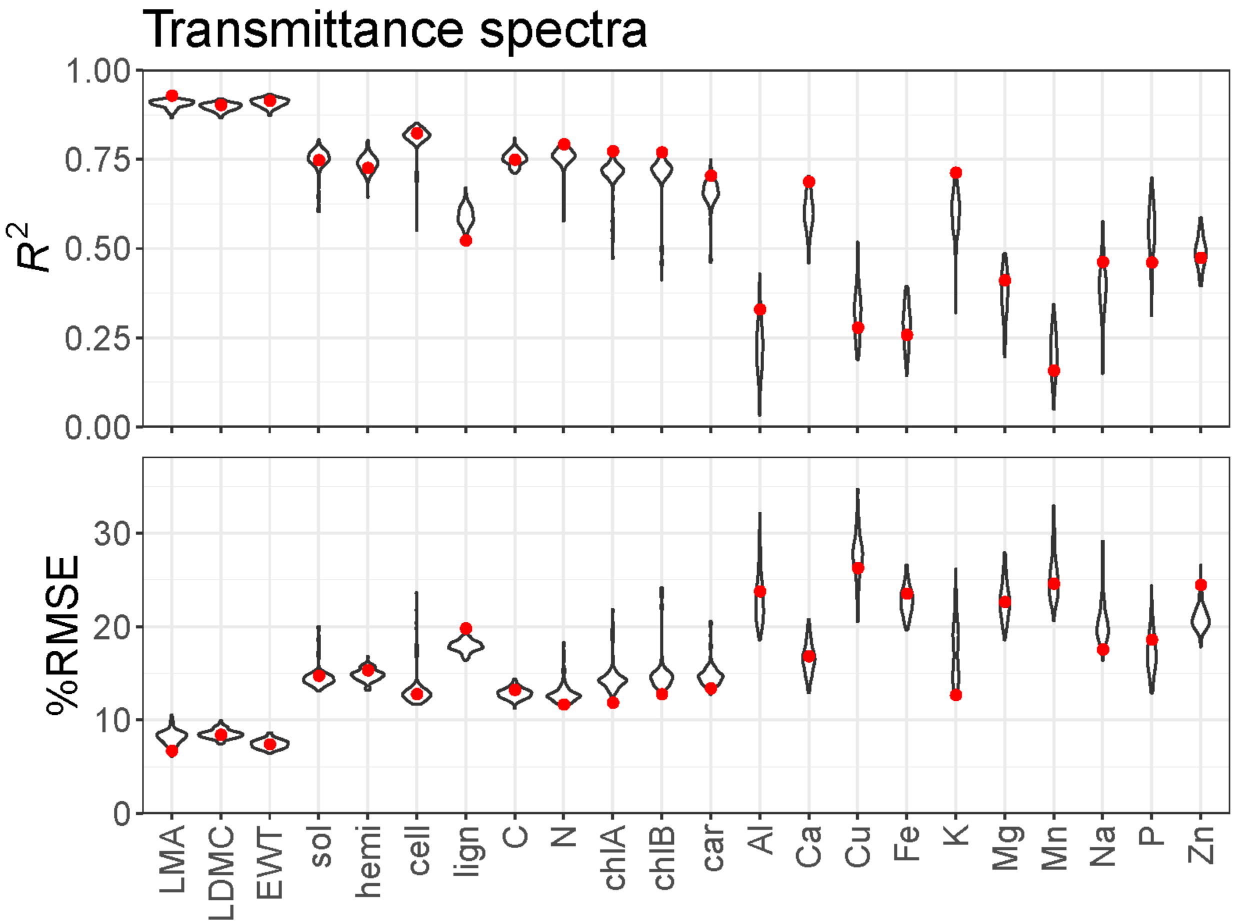

**Fig. S12:** The distributions of absorptance-based model performance statistics for each trait based on 100 jackknife resampling iterations from the calibration data set. The red dots show the $R^2$ and %RMSE from applying the ensemble of models to the internal validation subset. Abbreviations: sol = solubles, hemi = hemicellulose, cell = cellulose, lign = lignin, car = total carotenoids.

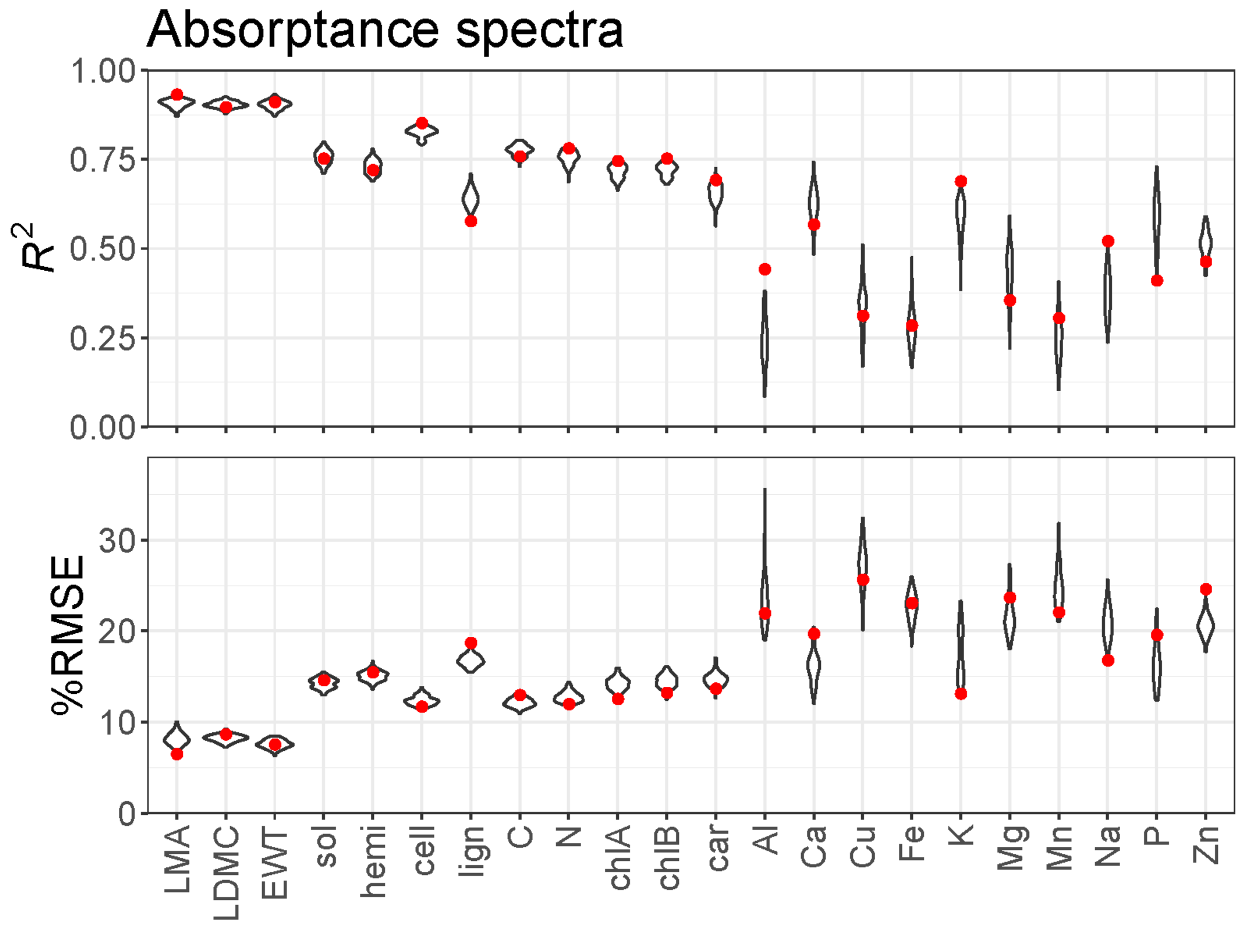

**Fig. S13:** The variable influence on projection (VIP) metric for reflectance-based models (Wold et al. 2001). The dashed line at 0.8 represents a heuristic threshold for band importance recommended by Burnett et al. (2021).

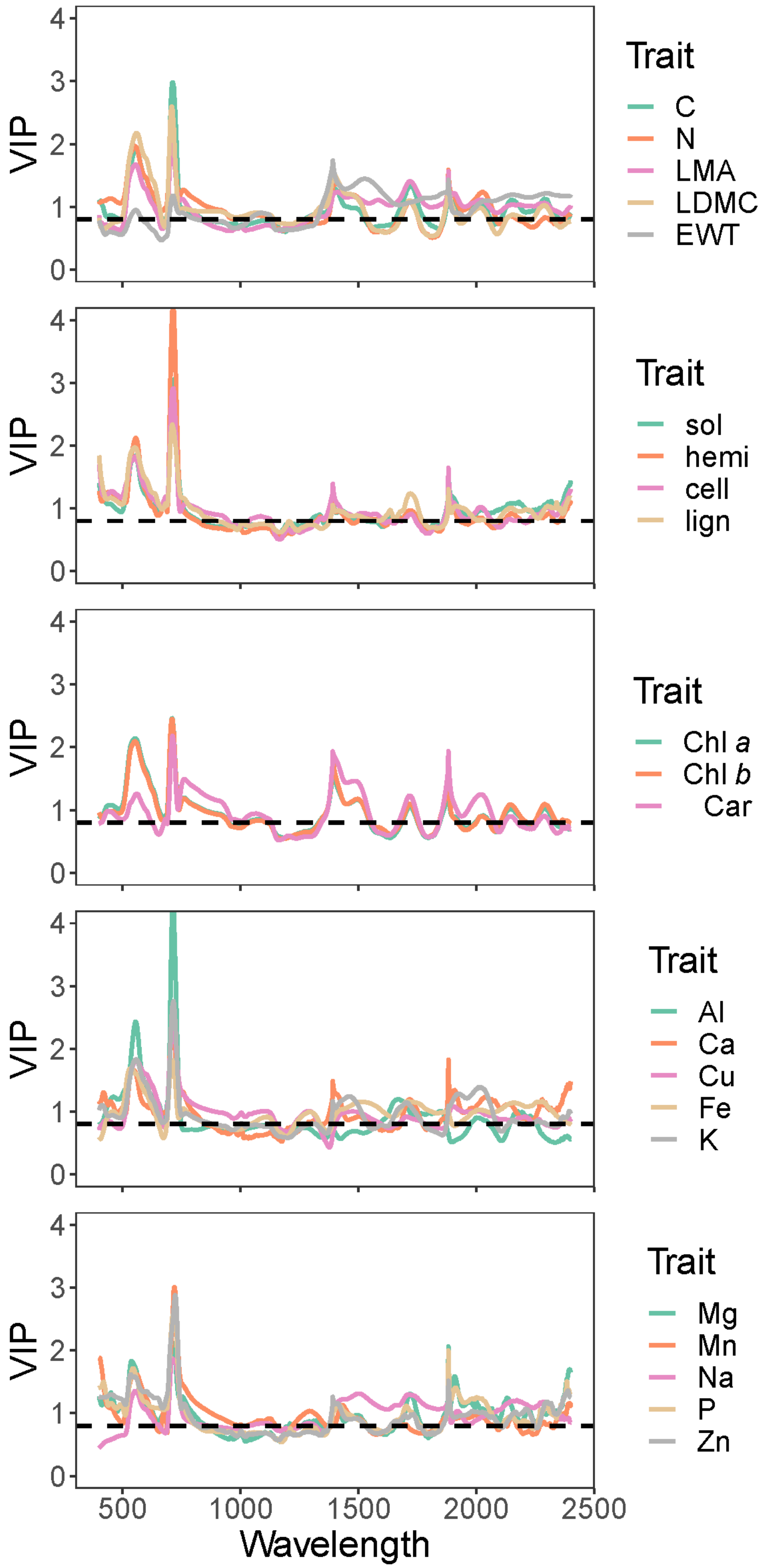

VIP
Wavelength
0
1
2
3
4
500
1000
1500
2000
2500
Trait
C
N
LMA
LDMC
EWT
Trait
sol
hemi
cell
lign
Trait
Chl *a*
Chl *b*
Car
Trait
Al
Ca
Cu
Fe
K
Trait
Mg
Mn
Na
P
Zn


290

**Fig. S14:** The variable influence on projection (VIP) metric for transmittance-based models (Wold et al. 2001). The dashed line at 0.8 represents a heuristic threshold for band importance recommended by Burnett et al. (2021).

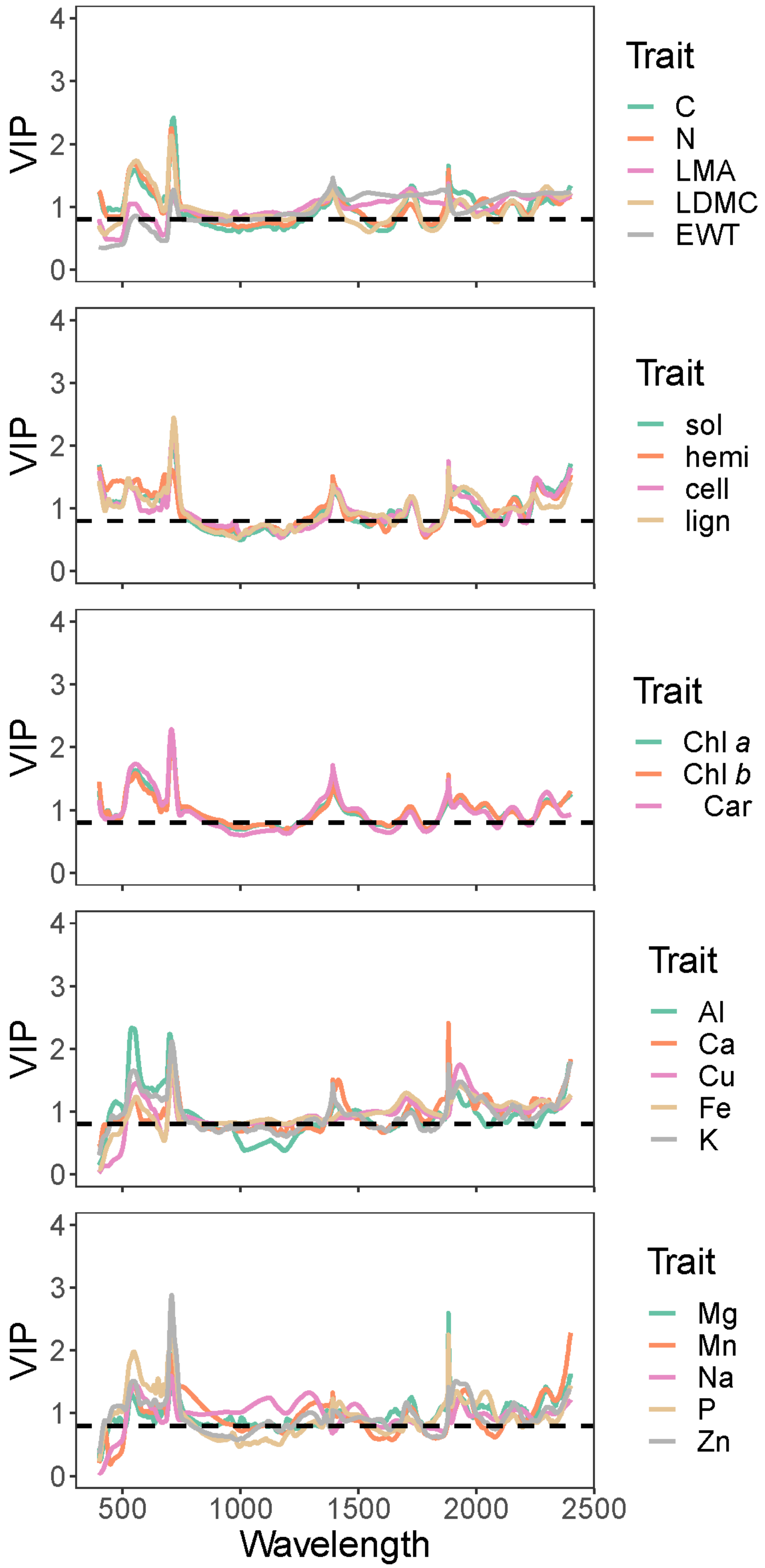
VIP
Trait
C
N
LMA
LDMC
EWT
sol
hemi
cell
lign
Chl a
Chl b
Car
Al
Ca
Cu
Fe
K
Mg
Mn
Na
P
Zn
0
1
2
3
4
500
1000
1500
2000
2500
Wavelength

**Fig. S15:** The variable influence on projection (VIP) metric for absorptance-based models (Wold et al. 2001). The dashed line at 0.8 represents a heuristic threshold for band importance recommended by Burnett et al. (2021).

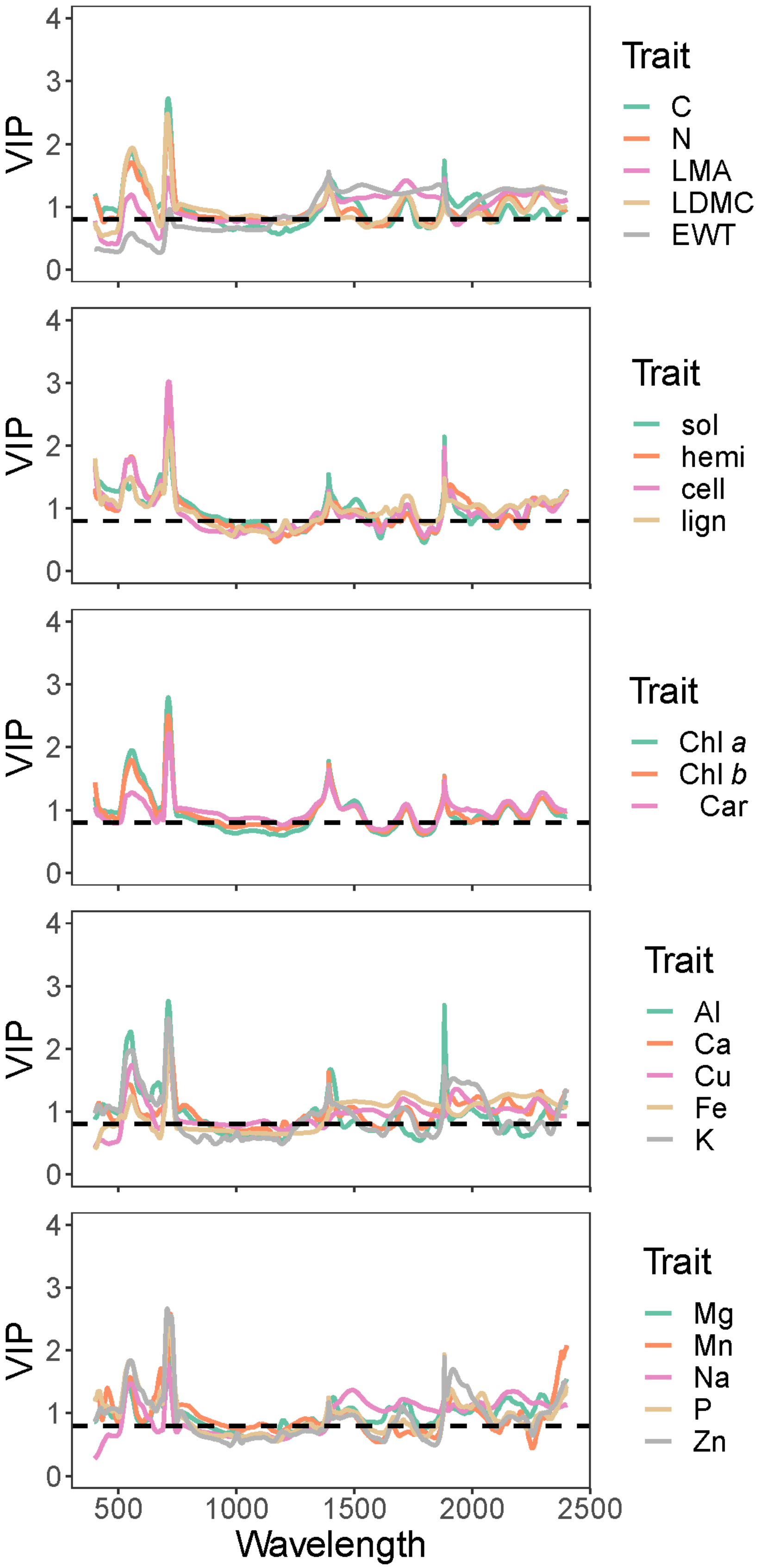

VIP
Trait
C
N
LMA
LDMC
EWT
sol
hemi
cell
lign
Chl a
Chl b
Car
Al
Ca
Cu
Fe
K
Mg
Mn
Na
P
Zn
Wavelength


298

**Fig. S16:** Plots of observations against reflectance-based PLSR predictions among internal validation data for various area-based leaf chemical traits. The black dashed line in each panel is the 1:1 line. Colored lines represent best-fit lines from OLS regression for each functional group. Error bars around each point represent 95% confidence intervals based on the ensemble of models produced in the 100× jackknife analysis.

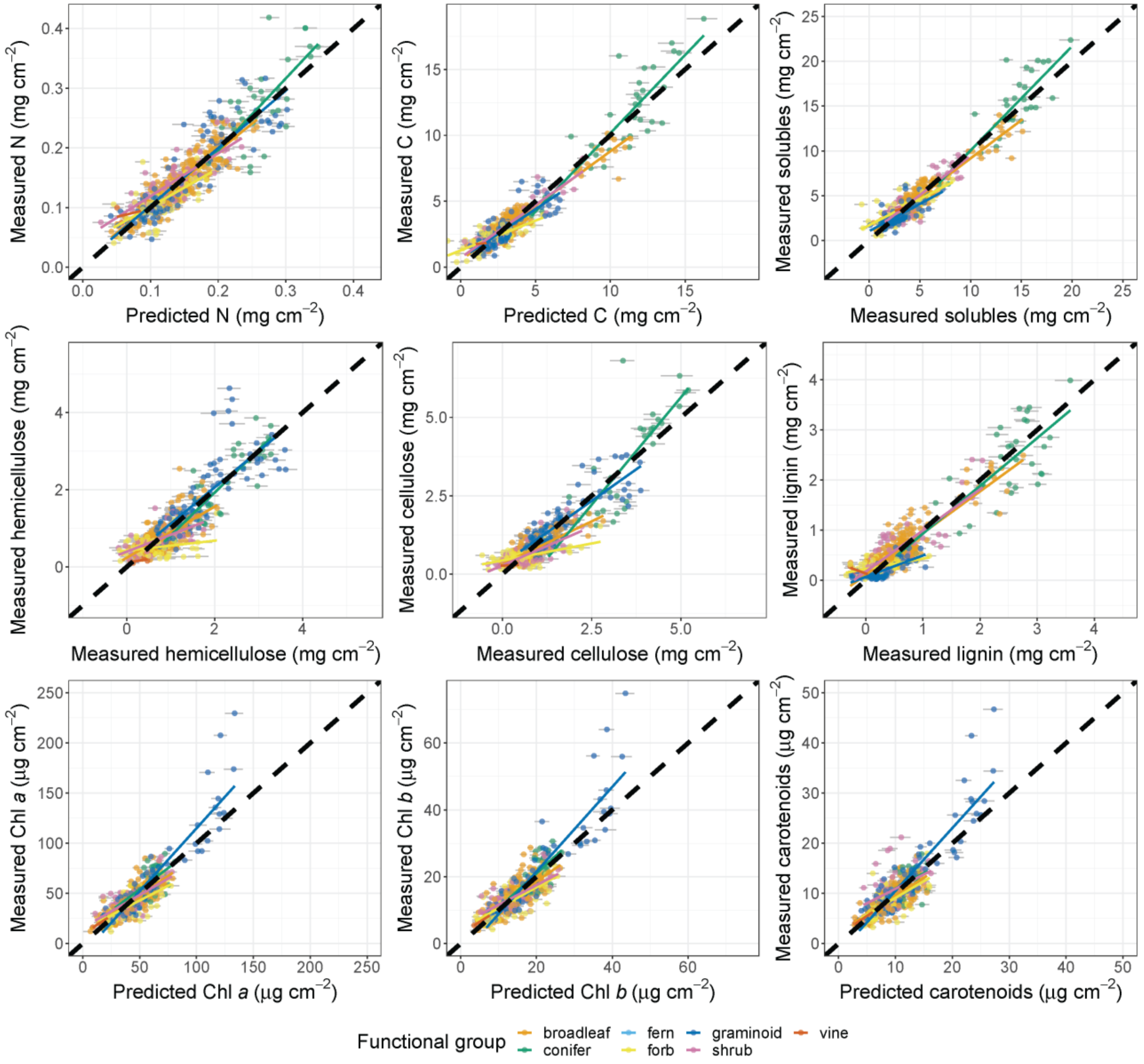

**Fig. S17:** Plots of observations against reflectance-based PLSR predictions among internal validation data for area-based contents of elements other than C and N. The black dashed line in each panel is the 1:1 line. Colored lines represent best-fit lines from OLS regression for each functional group. Error bars around each point represent 95% confidence intervals based on the ensemble of models produced in the 100× jackknife analysis.

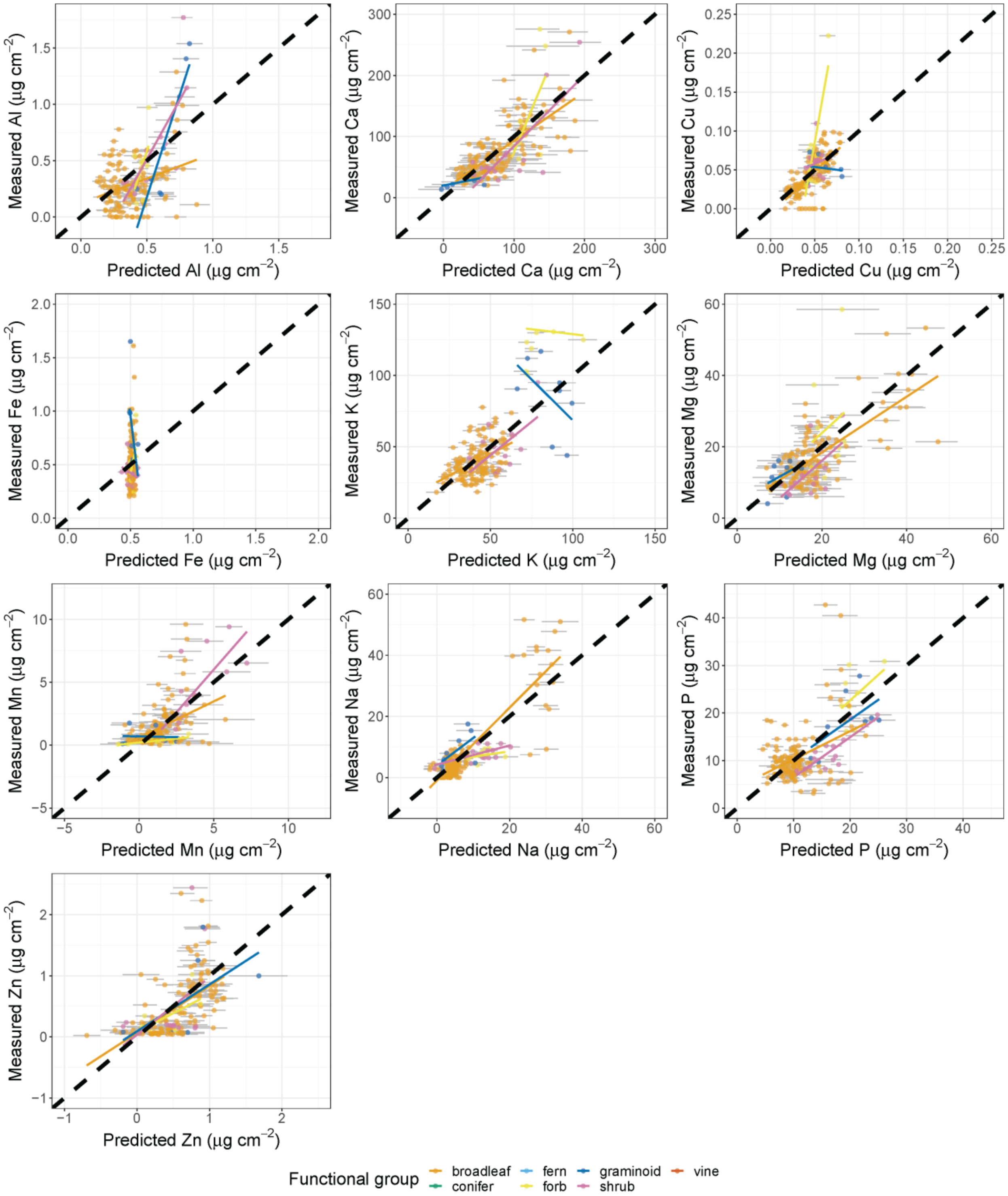
Measured Al (µg cm$^{-2}$)
Predicted Al (µg cm$^{-2}$)
Measured Ca (µg cm$^{-2}$)
Predicted Ca (µg cm$^{-2}$)
Measured Cu (µg cm$^{-2}$)
Predicted Cu (µg cm$^{-2}$)
Measured Fe (µg cm$^{-2}$)
Predicted Fe (µg cm$^{-2}$)
Measured K (µg cm$^{-2}$)
Predicted K (µg cm$^{-2}$)
Measured Mg (µg cm$^{-2}$)
Predicted Mg (µg cm$^{-2}$)
Measured Mn (µg cm$^{-2}$)
Predicted Mn (µg cm$^{-2}$)
Measured Na (µg cm$^{-2}$)
Predicted Na (µg cm$^{-2}$)
Measured P (µg cm$^{-2}$)
Predicted P (µg cm$^{-2}$)
Measured Zn (µg cm$^{-2}$)
Predicted Zn (µg cm$^{-2}$)
Functional group
broadleaf
conifer
fern
forb
graminoid
shrub
vine

**Fig. S18:** The distributions of reflectance-based model performance statistics for each area-based chemical trait based on 100 jackknife resampling iterations from the calibration data set. The red dots show the $R^2$ and %RMSE from applying the ensemble of models to the internal validation subset. Abbreviations: sol = solubles, hemi = hemicellulose, cell = cellulose, lign = lignin, car = total carotenoids.

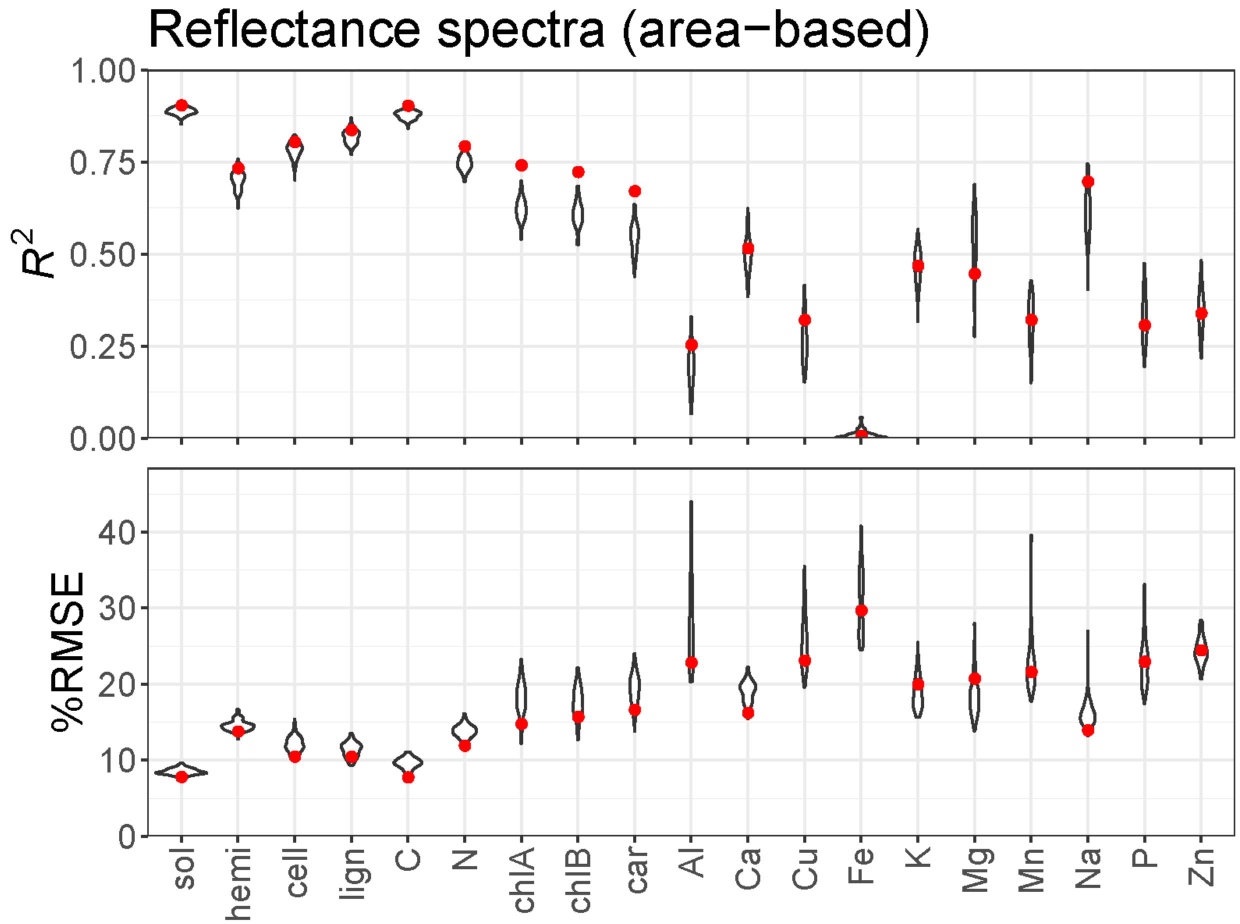

**Fig. S19:** The variable influence on projection (VIP) metric for reflectance-based models (Wold et al. 2001) for area-based chemical traits. The dashed line at 0.8 represents a heuristic threshold for band importance recommended by Burnett et al. (2021).

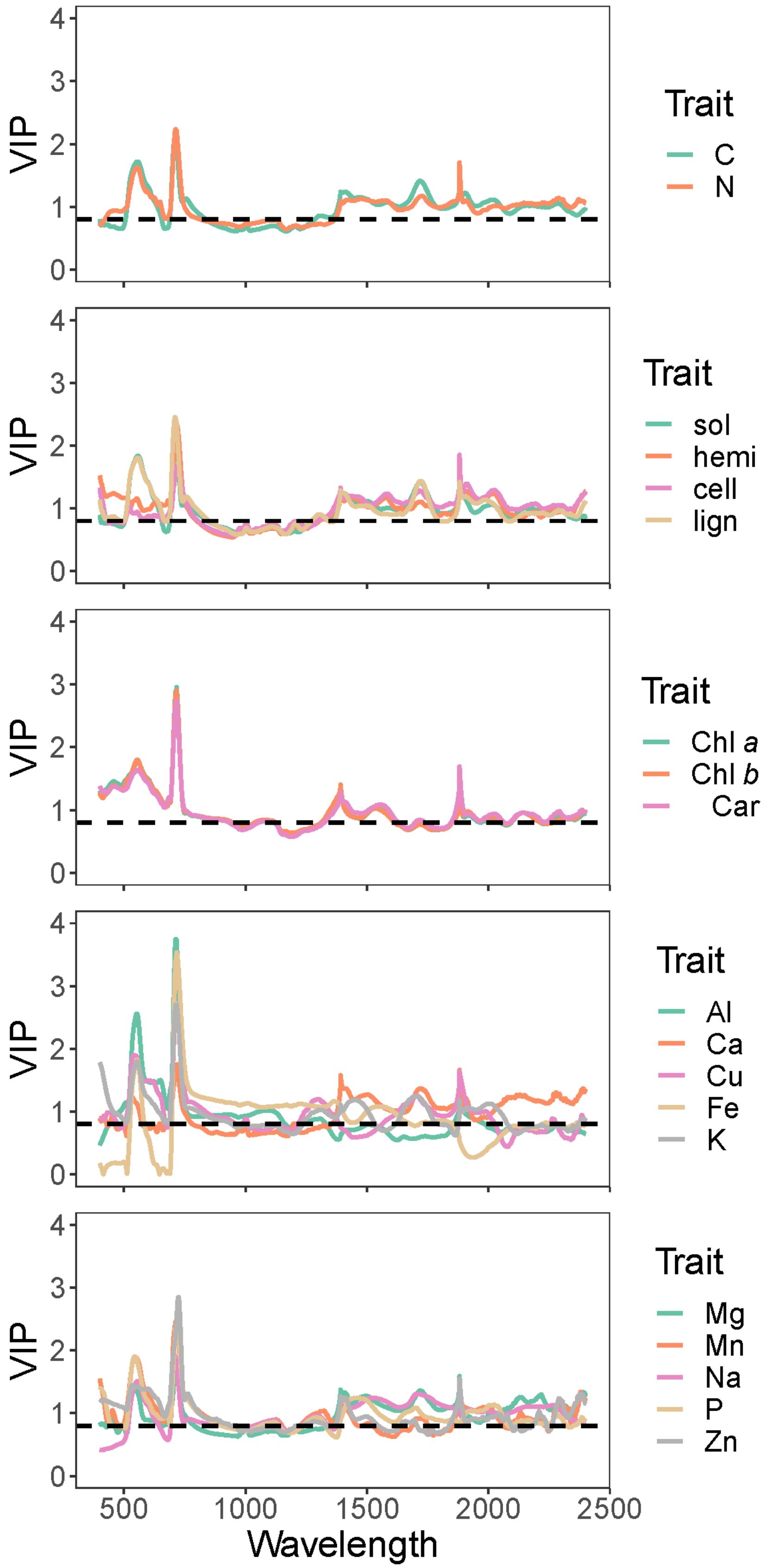
VIP
Trait
C
N
sol
hemi
cell
lign
Chl *a*
Chl *b*
Car
Al
Ca
Cu
Fe
K
Mg
Mn
Na
P
Zn
0
1
2
3
4
500
1000
1500
2000
2500
Wavelength

**Fig. S20:** Plots of observations against reflectance-based PLSR predictions among external validation data for elements other than C and N. The black dashed line is the 1:1 line. Error bars around each point represent 95% confidence intervals based on the ensemble of models produced in the 100× jackknife analysis. The colored line represents the best-fit line from OLS regression.

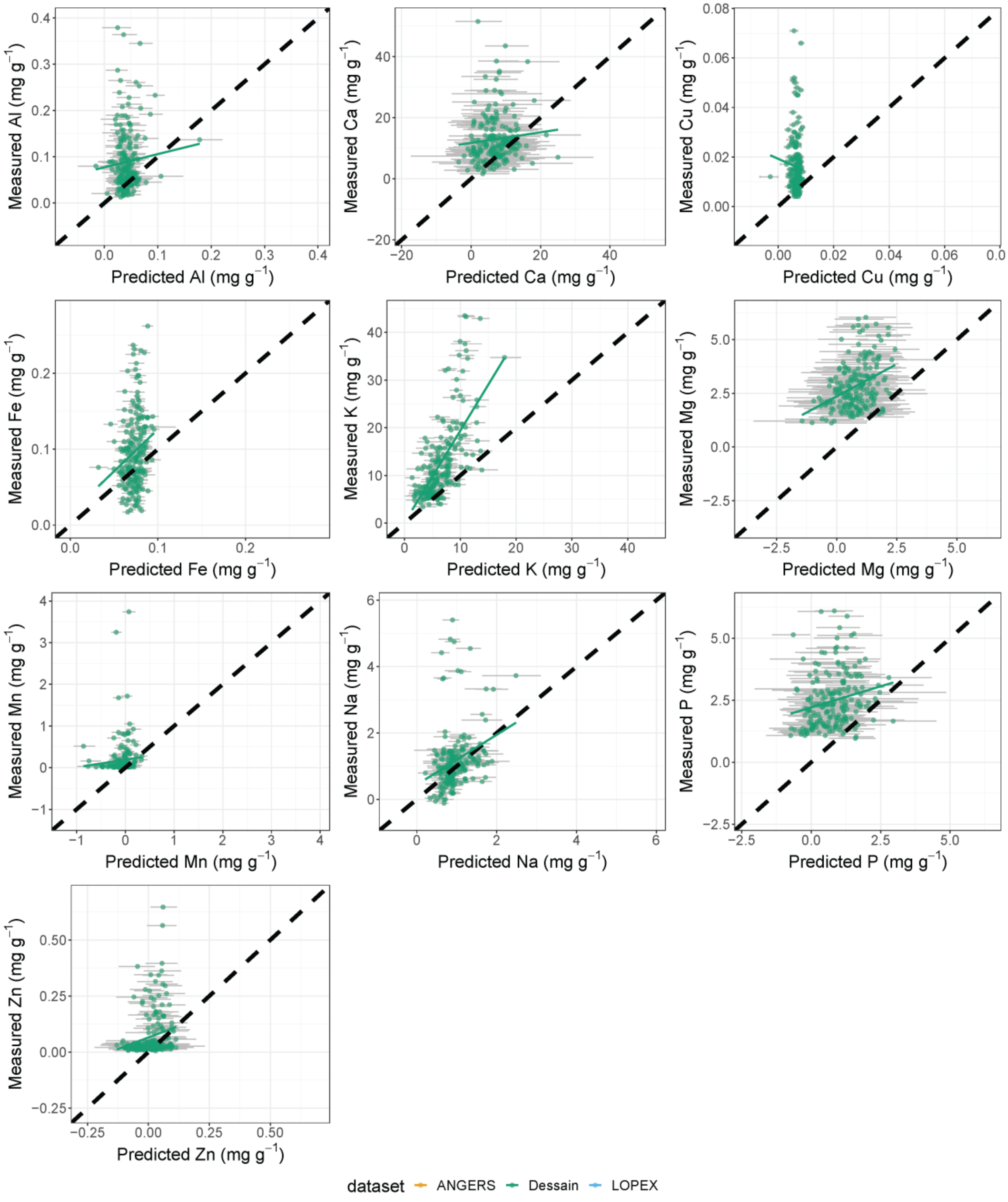

Measured Al (mg g−1)
Predicted Al (mg g−1)
Measured Ca (mg g−1)
Predicted Ca (mg g−1)
Measured Cu (mg g−1)
Predicted Cu (mg g−1)
Measured Fe (mg g−1)
Predicted Fe (mg g−1)
Measured K (mg g−1)
Predicted K (mg g−1)
Measured Mg (mg g−1)
Predicted Mg (mg g−1)
Measured Mn (mg g−1)
Predicted Mn (mg g−1)
Measured Na (mg g−1)
Predicted Na (mg g−1)
Measured P (mg g−1)
Predicted P (mg g−1)
Measured Zn (mg g−1)
Predicted Zn (mg g−1)
dataset
ANGERS
Dessain
LOPEX

**Supplementary tables**

**Table S1:** Summary statistics for internal validations of models predicting area-based leaf chemical traits from reflectance spectra. %RMSE is calculated as RMSE divided by the inner 95% trait range. Mass-proportionality is calculated as in Osnas et al. (2013) as the slope fitting the area-normalized trait vs. LMA, both $log_{10}$-transformed.

| | | | Area-based estimates | | | Mass-based estimates × LMA estimates | | |
|---|---|---|---|---|---|---|---|---|
| Trait | Mass-proportionality | # comps | RMSE | %RMSE | $R^2$ | RMSE | %RMSE | $R^2$ |
| N (mg cm$^{-2}$) | 0.591 | 20 | 0.0279 | 11.9 | 0.792 | 0.0459 | 19.5 | 0.542 |
| C (mg cm$^{-2}$) | 1.057 | 24 | 0.856 | 7.70 | 0.902 | 0.830 | 7.47 | 0.908 |
| Solubles (mg cm$^{-2}$) | 1.000 | 22 | 1.10 | 7.76 | 0.903 | 1.18 | 8.30 | 0.891 |
| Hemicellulose (mg cm$^{-2}$) | 0.834 | 22 | 0.404 | 13.7 | 0.733 | 0.454 | 15.4 | 0.685 |
| Cellulose (mg cm$^{-2}$) | 1.010 | 25 | 0.442 | 10.4 | 0.804 | 0.398 | 9.39 | 0.840 |
| Lignin (mg cm$^{-2}$) | 1.361 | 22 | 0.272 | 10.4 | 0.836 | 0.281 | 10.8 | 0.827 |
| Chl *a* (μg cm$^{-2}$) | 0.316 | 14 | 11.9 | 14.7 | 0.741 | 16.7 | 20.5 | 0.609 |
| Chl *b* (μg cm$^{-2}$) | 0.308 | 14 | 3.96 | 15.6 | 0.722 | 5.63 | 22.2 | 0.574 |
| Carotenoids (μg cm$^{-2}$) | 0.311 | 15 | 2.68 | 16.6 | 0.671 | 3.69 | 22.8 | 0.518 |
| Al (μg cm$^{-2}$) | 0.865 | 4 | 0.265 | 22.8 | 0.253 | 0.253 | 21.8 | 0.295 |
| Ca (μg cm$^{-2}$) | 0.435 | 17 | 35.7 | 16.2 | 0.515 | 41.3 | 18.7 | 0.366 |
| Cu (μg cm$^{-2}$) | -0.139 | 6 | 0.0223 | 23.1 | 0.321 | 0.0262 | 27.1 | 0.089 |
| Fe (μg cm$^{-2}$) | 0.215 | 1 | 0.230 | 29.6 | 0.006 | 0.234 | 30.2 | 0.021 |
| K (μg cm$^{-2}$) | 0.335 | 7 | 19.9 | 19.9 | 0.468 | 21.5 | 21.5 | 0.372 |
| Mg (μg cm$^{-2}$) | 0.544 | 15 | 6.97 | 20.7 | 0.446 | 7.69 | 22.8 | 0.389 |
| Mn (μg cm$^{-2}$) | 0.905 | 11 | 1.60 | 21.5 | 0.321 | 1.70 | 22.8 | 0.263 |
| Na (μg cm$^{-2}$) | 1.906 | 6 | 5.79 | 13.9 | 0.696 | 5.54 | 13.3 | 0.717 |
| P (μg cm$^{-2}$) | 0.421 | 6 | 5.51 | 22.9 | 0.307 | 6.13 | 25.5 | 0.238 |
| Zn (μg cm$^{-2}$) | -0.084 | 12 | 0.428 | 24.4 | 0.338 | 0.436 | 24.9 | 0.347 |

**Table S2:** Summary statistics for internal validations of models predicting leaf traits from reflectance (R), transmittance (T), and absorptance (A) spectra. %RMSE is calculated as RMSE divided by the inner 95% trait range.

| Trait | # comps | | | RMSE | | | %RMSE | | | $R^2$ | | |
|---|---|---|---|---|---|---|---|---|---|---|---|---|
| | R | T | A | R | T | A | R | T | A | R | T | A |
| LMA (kg $m^{-2}$) | 23 | 27 | 26 | 0.0175 | 0.0146 | 0.0141 | 8.04 | 6.69 | 6.49 | 0.892 | 0.928 | 0.930 |
| LDMC (mg $g^{-1}$) | 20 | 21 | 19 | 32.9 | 28.3 | 29.1 | 9.78 | 8.4 | 8.66 | 0.867 | 0.901 | 0.895 |
| EWT (mm) | 13 | 13 | 12 | 0.0334 | 0.0292 | 0.0298 | 8.46 | 7.38 | 7.54 | 0.885 | 0.913 | 0.910 |
| N (%) | 19 | 22 | 19 | 0.403 | 0.342 | 0.351 | 13.7 | 11.6 | 12.0 | 0.709 | 0.791 | 0.780 |
| C (%) | 20 | 25 | 23 | 1.38 | 1.38 | 1.35 | 13.2 | 13.2 | 13.0 | 0.747 | 0.748 | 0.757 |
| Solubles (%) | 23 | 24 | 25 | 6.59 | 6.41 | 6.35 | 15.1 | 14.7 | 14.6 | 0.733 | 0.747 | 0.751 |
| Hemicellulose (%) | 20 | 23 | 22 | 4.35 | 4.12 | 4.17 | 16.1 | 15.3 | 15.5 | 0.695 | 0.725 | 0.719 |
| Cellulose (%) | 25 | 26 | 26 | 2.74 | 2.76 | 2.53 | 12.7 | 12.8 | 11.7 | 0.826 | 0.822 | 0.851 |
| Lignin (%) | 20 | 23 | 22 | 2.82 | 2.93 | 2.77 | 19.1 | 19.8 | 18.7 | 0.558 | 0.522 | 0.576 |
| Chl *a* (mg $g^{-1}$) | 13 | 20 | 16 | 2.07 | 1.77 | 1.87 | 13.8 | 11.8 | 12.5 | 0.689 | 0.772 | 0.744 |
| Chl *b* (mg $g^{-1}$) | 14 | 23 | 18 | 0.716 | 0.607 | 0.63 | 15.0 | 12.7 | 13.2 | 0.679 | 0.769 | 0.751 |
| Carotenoids (mg $g^{-1}$) | 12 | 16 | 13 | 0.434 | 0.396 | 0.404 | 14.7 | 13.4 | 13.7 | 0.644 | 0.703 | 0.691 |
| Al (mg $g^{-1}$) | 6 | 9 | 10 | 0.0293 | 0.0283 | 0.0261 | 24.6 | 23.7 | 21.9 | 0.304 | 0.328 | 0.441 |
| Ca (mg $g^{-1}$) | 17 | 22 | 17 | 4.03 | 3.32 | 3.88 | 20.4 | 16.8 | 19.7 | 0.541 | 0.686 | 0.566 |
| Cu (mg $g^{-1}$) | 5 | 4 | 5 | 0.00311 | 0.00315 | 0.00307 | 25.9 | 26.2 | 25.6 | 0.295 | 0.278 | 0.311 |
| Fe (mg $g^{-1}$) | 4 | 6 | 5 | 0.0295 | 0.0301 | 0.0295 | 23.0 | 23.5 | 23.0 | 0.286 | 0.257 | 0.283 |
| K (mg $g^{-1}$) | 10 | 18 | 18 | 2.32 | 1.87 | 1.94 | 15.7 | 12.7 | 13.1 | 0.55 | 0.712 | 0.688 |
| Mg (mg $g^{-1}$) | 18 | 17 | 16 | 0.714 | 0.708 | 0.74 | 22.8 | 22.6 | 23.6 | 0.394 | 0.410 | 0.354 |
| Mn (mg $g^{-1}$) | 10 | 8 | 11 | 0.164 | 0.177 | 0.158 | 22.8 | 24.6 | 22.0 | 0.254 | 0.157 | 0.304 |
| Na (mg $g^{-1}$) | 6 | 4 | 5 | 0.451 | 0.472 | 0.452 | 16.7 | 17.5 | 16.8 | 0.519 | 0.461 | 0.520 |
| P (mg $g^{-1}$) | 15 | 20 | 20 | 0.607 | 0.550 | 0.579 | 20.5 | 18.6 | 19.6 | 0.313 | 0.460 | 0.410 |
| Zn (mg $g^{-1}$) | 17 | 12 | 13 | 0.0486 | 0.0491 | 0.0493 | 24.2 | 24.4 | 24.5 | 0.487 | 0.473 | 0.462 |

**Table S3:** Summary statistics for the performance of models calibrated on continuum-removed CABO data and applied to external validation datasets.

| | Dessain | | | LOPEX | | | ANGERS | | |
|---|---|---|---|---|---|---|---|---|---|
| | $R^2$ | RMSE | %RMSE | $R^2$ | RMSE | %RMSE | $R^2$ | RMSE | %RMSE |
| LMA (kg $m^{-2}$) | 0.364 | 0.0907 | 131 | 0.435 | 0.0976 | 98.3 | 0.444 | 0.130 | 111 |
| LDMC (mg $g^{-1}$) | 0.840 | 33.1 | 11.1 | 0.893 | 55.2 | 14.3 | | | |
| EWT (mm) | 0.617 | 0.0477 | 29.2 | 0.861 | 0.0661 | 25.8 | 0.774 | 0.0751 | 39.3 |
| N (%) | 0.409 | 0.741 | 24.1 | 0.505 | 0.837 | 19.8 | | | |
| C (%) | 0.447 | 3.21 | 45.5 | 0.163 | 3.81 | 34.5 | | | |
| Solubles (%) | 0.222 | 10.9 | 26.8 | | | | | | |
| Hemicellulose (%) | 0.263 | 5.98 | 22.3 | | | | | | |
| Cellulose (%) | 0.324 | 3.86 | 25.2 | 0.243 | 9.05 | 38.3 | | | |
| Lignin (%) | 0.140 | 4.44 | 28.4 | 0.158 | 4.98 | 27.9 | | | |
| Chl *a* (mg $g^{-1}$) | 0.354 | 2.38 | 22.1 | 0.411 | 3.53 | 36.4 | 0.473 | 4.06 | 31.7 |
| Chl *b* (mg $g^{-1}$) | 0.369 | 0.842 | 25.2 | 0.431 | 1.23 | 31.1 | 0.347 | 1.99 | 43.1 |
| Carotenoids (mg $g^{-1}$) | 0.213 | 0.517 | 23.9 | 0.307 | 0.64 | 22.8 | 0.376 | 0.934 | 25.6 |
| Al (mg $g^{-1}$) | 0.001 | 0.0763 | 31.5 | | | | | | |
| Ca (mg $g^{-1}$) | 0.011 | 18.1 | 57.2 | | | | | | |
| Cu (mg $g^{-1}$) | 0.001 | 0.0144 | 33.4 | | | | | | |
| Fe (mg $g^{-1}$) | 0.000 | 0.0594 | 29.8 | | | | | | |
| K (mg $g^{-1}$) | 0.322 | 8.04 | 25.5 | | | | | | |
| Mg (mg $g^{-1}$) | 0.107 | 2.89 | 67.3 | | | | | | |
| Mn (mg $g^{-1}$) | 0.024 | 0.568 | 61.5 | | | | | | |
| Na (mg $g^{-1}$) | 0.062 | 0.903 | 23.2 | | | | | | |
| P (mg $g^{-1}$) | 0.041 | 3.30 | 82 | | | | | | |
| Zn (mg $g^{-1}$) | 0.054 | 0.107 | 31.8 | | | | | | |

**Table S4:** Summary statistics for the performance of models calibrated on brightness-normalized CABO data and applied to external validation datasets.

| | Dessain | | | LOPEX | | | ANGERS | | |
|---|---|---|---|---|---|---|---|---|---|
| | $R^2$ | RMSE | %RMSE | $R^2$ | RMSE | %RMSE | $R^2$ | RMSE | %RMSE |
| LMA (kg $m^{-2}$) | 0.658 | 0.0700 | 101 | 0.735 | 0.0879 | 88.6 | 0.540 | 0.127 | 108 |
| LDMC (mg $g^{-1}$) | 0.812 | 41.7 | 14 | 0.881 | 45.7 | 11.8 | | | |
| EWT (mm) | 0.614 | 0.0422 | 25.8 | 0.905 | 0.0619 | 24.1 | 0.797 | 0.0751 | 39.2 |
| N (%) | 0.475 | 0.659 | 21.4 | 0.541 | 0.918 | 21.8 | | | |
| C (%) | 0.413 | 1.91 | 27.1 | 0.114 | 2.85 | 25.8 | | | |
| Solubles (%) | 0.306 | 12.2 | 30.1 | | | | | | |
| Hemicellulose (%) | 0.353 | 4.91 | 18.3 | | | | | | |
| Cellulose (%) | 0.307 | 7.41 | 48.3 | 0.358 | 5.37 | 22.7 | | | |
| Lignin (%) | 0.187 | 4.59 | 29.3 | 0.153 | 4.89 | 27.4 | | | |
| Chl *a* (mg $g^{-1}$) | 0.333 | 2.75 | 25.5 | 0.392 | 3.44 | 35.5 | 0.548 | 2.8 | 21.9 |
| Chl *b* (mg $g^{-1}$) | 0.342 | 0.964 | 28.9 | 0.451 | 1.17 | 29.6 | 0.426 | 1.45 | 31.4 |
| Carotenoids (mg $g^{-1}$) | 0.193 | 0.568 | 26.3 | 0.270 | 0.656 | 23.4 | 0.491 | 1.01 | 27.7 |
| Al (mg $g^{-1}$) | 0.003 | 0.0742 | 30.6 | | | | | | |
| Ca (mg $g^{-1}$) | 0.005 | 11.7 | 37 | | | | | | |
| Cu (mg $g^{-1}$) | 0.001 | 0.0145 | 33.4 | | | | | | |
| Fe (mg $g^{-1}$) | 0.046 | 0.0526 | 26.4 | | | | | | |
| K (mg $g^{-1}$) | 0.468 | 9.02 | 28.6 | | | | | | |
| Mg (mg $g^{-1}$) | 0.123 | 2.37 | 55.1 | | | | | | |
| Mn (mg $g^{-1}$) | 0.004 | 0.503 | 54.5 | | | | | | |
| Na (mg $g^{-1}$) | 0.084 | 0.907 | 23.3 | | | | | | |
| P (mg $g^{-1}$) | 0.063 | 2.33 | 58 | | | | | | |
| Zn (mg $g^{-1}$) | 0.037 | 0.107 | 31.7 | | | | | | |

**Table S5:** Summary statistics from model transferability analyses among functional groups. Each functional group in turn, other than ferns and vines, was left out of the dataset used to train a PLSR model for each trait using 10-fold cross-validation. The functional group left out (column names) was then used for validation.

| | Broadleaf | | | Conifer | | | Forb | | | Graminoid | | | Shrub | | |
|---|---|---|---|---|---|---|---|---|---|---|---|---|---|---|---|
| | $R^2$ | RMSE | %RMSE | $R^2$ | RMSE | %RMSE | $R^2$ | RMSE | %RMSE | $R^2$ | RMSE | %RMSE | $R^2$ | RMSE | %RMSE |
| LMA (kg $m^{-2}$) | 0.764 | 0.0249 | 12.0 | 0.413 | 0.104 | 50.3 | 0.300 | 0.0278 | 13.4 | 0.552 | 0.0252 | 12.2 | 0.879 | 0.0230 | 11.1 |
| LDMC (mg $g^{-1}$) | 0.668 | 37.6 | 10.7 | 0.517 | 40.2 | 11.5 | 0.814 | 39.0 | 11.1 | 0.675 | 57.2 | 16.3 | 0.832 | 33.7 | 9.61 |
| EWT (mm) | 0.453 | 0.0362 | 9.2 | 0.575 | 0.104 | 26.4 | 0.572 | 0.0555 | 14.1 | 0.906 | 0.0652 | 16.6 | 0.600 | 0.0329 | 8.34 |
| N (%) | 0.599 | 0.592 | 21.3 | 0.577 | 0.418 | 15.1 | 0.393 | 0.673 | 24.3 | 0.409 | 0.566 | 20.4 | 0.734 | 0.584 | 21.1 |
| C (%) | 0.468 | 1.54 | 14.5 | 0.239 | 1.88 | 17.7 | 0.035 | 2.13 | 20.0 | 0.089 | 2.16 | 20.3 | 0.746 | 2.28 | 21.4 |
| Solubles (%) | 0.357 | 8.15 | 18.3 | 0.266 | 9.67 | 21.7 | 0.108 | 15.4 | 34.6 | 0.086 | 22.4 | 50.2 | 0.292 | 8.99 | 20.2 |
| Hemicellulose (%) | 0.262 | 4.43 | 16.0 | 0.084 | 5.51 | 19.9 | 0.041 | 9.37 | 33.8 | 0.067 | 16.0 | 57.8 | 0.255 | 5.68 | 20.5 |
| Cellulose (%) | 0.285 | 4.08 | 18.4 | 0.358 | 5.64 | 25.5 | 0.090 | 6.74 | 30.5 | 0.061 | 11.6 | 52.4 | 0.002 | 4.63 | 20.9 |
| Lignin (%) | 0.182 | 3.44 | 22.4 | 0.358 | 2.76 | 18.0 | 0.116 | 3.60 | 23.4 | 0.034 | 5.46 | 35.6 | 0.466 | 4.06 | 26.4 |
| Chl *a* (mg $g^{-1}$) | 0.667 | 2.31 | 16.5 | 0.535 | 1.60 | 11.5 | 0.210 | 4.63 | 33.1 | 0.316 | 2.88 | 20.6 | 0.750 | 2.41 | 17.2 |
| Chl *b* (mg $g^{-1}$) | 0.668 | 0.690 | 14.5 | 0.481 | 0.620 | 13.1 | 0.157 | 1.78 | 37.5 | 0.332 | 0.958 | 20.2 | 0.737 | 0.736 | 15.5 |
| Carotenoids (mg $g^{-1}$) | 0.629 | 0.478 | 16.5 | 0.406 | 0.358 | 12.3 | 0.196 | 0.841 | 28.9 | 0.187 | 0.620 | 21.3 | 0.566 | 0.634 | 21.8 |
